# Dynamic binning peak detection and assessment of various lipidomics liquid chromatography-mass spectrometry pre-processing platforms

**DOI:** 10.1101/2020.10.10.334342

**Authors:** Xiaodong Feng, Wenxuan Zhang, Folkert Kuipers, Ido Kema, Andrei Barcaru, Péter Horvatovich

**Affiliations:** Department of Laboratory Medicine, University Medical Center Groningen, Hanzeplein 1, 9713 GZ Groningen, The Netherlands; Department of Pediatrics, University Medical Center Groningen, Hanzeplein 1, 9713 GZ Groningen, The Netherlands; Department of Analytical Biochemistry, University of Groningen, Antonius Deusinglaan 1, 9713 AV Groningen, The Netherlands

**Keywords:** Dynamic binning, peak detection, lipidomics, LC-MS pre-processing

## Abstract

Liquid chromatography-mass spectrometry (LC-MS) based lipidomics generate a large dataset, which requires high-performance data pre-processing tools for their interpretation such as XCMS, mzMine and Progenesis. These pre-processing tools rely heavily on accurate peak detection, which depends on setting the peak detection mass tolerance (PDMT) properly. The PDMT is usually set with a fixed value in either ppm or Da units. However, this fixed value may result in duplicates or missed peak detection. Therefore, we developed the dynamic binning method for accurate peak detection, which takes into account the peak broadening described by well-known physics laws of ion separation and set dynamically the value of PDMT as a function of m/z. Namely, in our method, the PDMT is proportional to 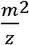 for FTICR, to 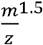 for Orbitrap, to *m/z* for Q-TOF and is a constant for Quadrupole mass analyzer, respectively. The dynamic binning method was implemented in XCMS [1,2] and the adopted source code is available in the Appendix. Our further goal was to compare the performance of different lipidomics pre-processing tools to find differential compounds. We have generated set samples with 43 lipids internal standards differentially spiked to aliquots of one human plasma lipid sample using Orbitrap LC-MS/MS. The performance of the various pipelines using aligned parameter sets was quantified by a quality score system which reflects the ability of a pre-processing pipeline to detect differential peaks spiked at various concentration levels. The quality score indicates that the dynamic binning method improves the performance of XCMS (maximum p-value 9.8·10^−3^ of two-sample Wilcoxon test). The modified XCMS software was further compared with mzMine and Progenesis. The results showed that modified XCMS and Progenesis had a similarly good performance in the aspect of finding differential compounds. In addition, Progenesis shows lower variability as indicated by lower CVs, followed by XCMS and mzMine. The lower variability of Progenesis improve the quantification, however, provide an incorrect quantification abundance order of spiked-in internal standards.

## 1. INTRODUCTION

In lipidomics, liquid chromatography-mass spectrometry (LC-MS) is commonly used quantitative profiling approach because of the high separation efficiency of LC and the large dynamic measurement range, the specificity and sensitivity of MS.[3,4] The large amounts of complex LC-MS data generated in typical experiment requires accurate quantitative pre-processing and identification of thousands of lipid species present in complex biological samples. Consequently, accurate data processing has become a major challenge in lipidomics.[5]

There is a significant amount of research directed to the development of LC-MS(/MS) data pre-processing tools. Some are designed primarily for lipidomics while others are more general LC-MS(/MS) pre-processing tools. To name a few, commercial tools such as *Progenesis*[6] developed by *Nonlinear Dynamics* and open source tools such as *mzMine*,[7,8] *XCMS*,[1,2] *KniMet*,[9] *OpenMS*,[10] *metaX*,[11] *LipidMatch*,[12] *CAMERA*,[13] *MetaboAnalystR*,[14] *MetFrag*;[15] and online tools like *XCMS online*,[16] *MetaboAnalyst*,[17] *PiMP my metabolome*,[18] *Workflow4Metabolomics*,[19] *MetFrag online*[15] are available in the field.

The above data processing tools typically contain the following processing modules[20]: data transformation such as resampling and smoothing, peak detection, retention time correction, the grouping of peaks across different samples, filling in missing data, annotation of individual and matched isotope peak clusters with potential metabolite identity, normalization of the quantification values and differential statistical analysis with univariate and multivariate methods or using other type of statistical analysis (Fig. A.1). Among them, peak detection is one of the most important step in an LC-MS(/MS) preprocessing workflows. Thus it is important to define an algorithm capable to distinguish between the irrelevant signal of chemical or electronic noise and a signal of a compound. The signal of a compound is usually represented as a 2-dimensional shaped Gaussian peak (Fig. A.1b) and peaks corresponding to different stable isotopes of the compound at given charge state form isotope cluster (Fig. A.1d).

Extracted ion chromatogram (EIC) - based peak detection algorithms typically consist of EIC construction and chromatographic peak detection from the constructed EICs. *XCMS*,[1] which was developed in 2006 to facilitate LC-MS data pre-processing, and is one of the most widely used tool for peak detection in metabolomics LC-MS datasets.[2],[21] It includes three EIC based peak detection algorithms suitable for LC-MS(/MS) pre-processing: *matchedFilter*, *centWave*,[22] and *massifquant*.[23] Among them, *centWave* is the most frequently used peak detection algorithm.

Despite the intensive efforts to develop and benchmark accurate peak detection, many currently used peak detection algorithms needs to improve in peak detection and quantification performance to avoid to report inaccurate quantification and/or false detection of compound[21] (i.e. false positives) and/or the inability to detect existing generally low abundant compounds[24] (i.e. false negatives) in LC-MS(/MS) data. False-positive and false-negative results may be due to the incorrect parameterization of the employed algorithm. Setting a proper mass tolerance is important for EIC construction (i.e. binning) used for peak detection. Although there are guidelines for setting the proper mass tolerance,[20] the recommended mass tolerance is not always suitable for specific instrumental settings and a specific m/z range. For example, a range of 5-15 ppm for Orbitrap data results in a too broad and overgeneralized range, which may result in mixing signal of multiple compounds. The mass range (thus peak width in *mz* dimension) generally increase in the function of the *mz* in case of high-resolution mass analyzers such as TOF, FTICR and Orbitrap instruments. (Note: for a clearer notation, we use *mz* across the entire article to represent *m/z*). Thus, *Mayers et al*. recommended using the mass tolerance units in Da (i.e. Daltons) instead of ppm (i.e. parts per million), as the mass range variation is smaller in Da units.[25] However, even if the mass tolerance is specified in Da, setting a fixed mass tolerance improperly may still result in duplicates or missed peak detection. Therefore, we applied a dynamic adjustment of mass tolerance to construct EICs to improve *centWave*’s[26] peak detection performance. In the original form of the *centWave* algorithm, mass tolerance is set in ppm and is thus increasing proportionally with *mz*. This relationship is correct for TOF mass analyzer, while is not optimal for Orbitrap mass analyzer, where the peak width and *mz* in mass spectra is proportional to 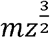.[27] Our dynamic binning method of constructing EICs takes into account the correct 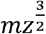 proportional peak broadening in *mz* for Orbitrap data in the peak detection step, which lead to more precise quantification.

LC-MS(/MS) pre-processing workflows differ in algorithmic design and involve many different parameters. It is difficult for a user to determine the correct parameters for peak detection to analyze a particular data. An objective evaluation of lipidomic LC-MS(/MS) data pre-processing workflows and used parameter set is important to compare the performance of different pipelines.[21,28] In 2012, *Hoekman et al*.[29] introduced a scoring method to compare the performances of different quantitative LC-MS/MS pre-processing workflows. The score quantifies the capacity of a pre-processing pipeline, with a given set of parameters, to detect differentially spiked compounds in sample that has the same composition i.e. consist of the same aliquot of one biological (background) sample. In this dataset the spiked compounds are not present in the background sample. We extended this scoring system to determine the distribution of scores which in turn allows to compare pipelines performance using non-parametric significance test. The modified scoring strategy was applied in this study to compare the performance of *centWave* peak picking with fixed and dynamic EIC construction tolerance in *XCMS*, as well as mzMine and Progenesis. Finally, we compared the performance of these LC-MS pre-processing workflows after optimizing the parameters for each workflow, to make them comparable as much as possible, despite the algorithmic differences.

## 2. THEORY

Fig. A.1 shows the most important aspects of the peak detection process, which consist of the following steps: EIC construction, peak detection, isotope pattern identification, adduct ion, isomer and feature detection.

### 2.1. Setting proper mass tolerance for accurate EIC construction

Peak detection provides crucial information that is required for accurate compound identification and quantification, and accurate construction of EIC is fundamental for accurate peak detection. Setting proper mass tolerance is important for a proper selection of the ions of one isotope of a compound that forms the EIC. Fig. A.1 shows how mass tolerance is related to the width of the selected rectangle. Although there is a detailed guideline for setting the proper mass tolerance in XCMS,[20] the mass tolerance is highly dependent on the *mz* and the type of mass analyser. As shown in Fig. 1, *compound 1* is located at a low *mz* of 376.3950 Da, which has a small mass peak width, around 0.003462 Da (9.2 ppm), while *compound 2* is located at high *mz* of 1428.1220 Da, which has a larger uncertainty and thus has a mass peak width around 0.025594 Da (18.0 ppm). Thus, a small value of mass tolerance may not be sufficient to cover the uncertainty of ions *mz* fluctuation in higher *mz*, resulting in missing intensity ions from peaks and to underestimation of peak quantity and may results in peak splitting. If a larger value is selected for a mass tolerance, merging of mass traces that are close can occur in lower *mz* leading to missing peaks and quantitative values of mixed peaks. Typically, this uncertainty (or variability) may result from two sources: the mass fluctuation (MF) and mass dispersion (MD). MF is fluctuation of the peak maxima in the mass spectrum, which can be observed between different subsequent MS1 mass spectra. Fig. A.2 and A.3 show MF exists in both low and high *mz*. This term sometimes is called mass accuracy by the vendor. For a well-calibrated Orbitrap, the MF should be less than 1 ppm.[30] MD can be estimated according to the relationship between the mass peak width and *mz*. MD is proportional to *mz*^*2*^ for FTICR, to *mz*^*1.5*^ for Orbitrap, to *mz* for Q-TOF and a constant value for Quadrupole.[27]

**Fig. 1.**
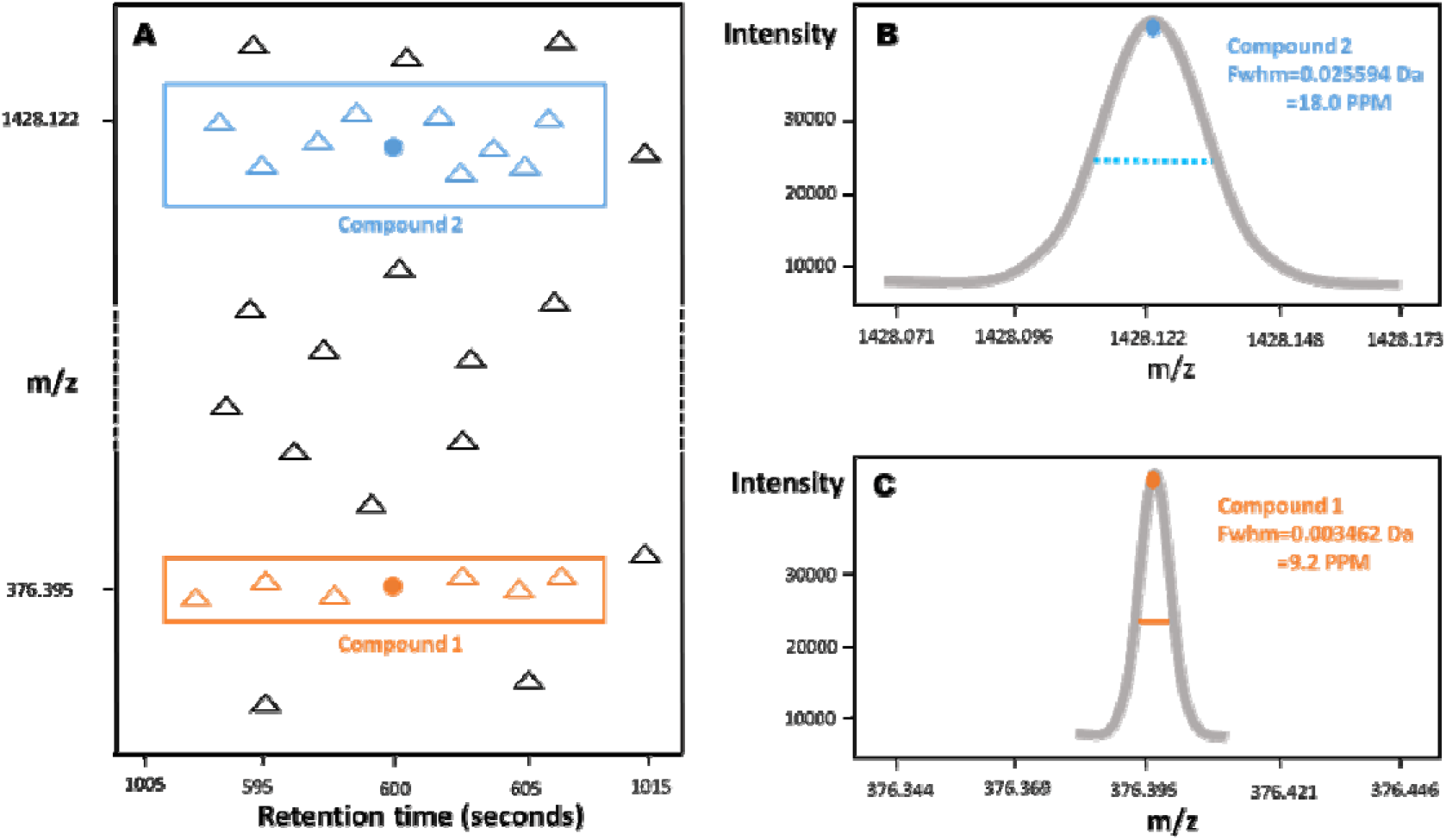
Scheme showing main aspects of the dynamic binning method. Compound 1 is located at a low *mz* of 376.3950 Da, which has a small mass peak width, around 0.003462 Da (9.2 ppm), while compound 2 is located at high *mz* of 1428.1220 Da, which has a larger peak width around 0.025594 Da (18.0 ppm). Dynamic binning is adopting the threshold tolerance of EIC construction in function of *mz* using equation derived from ion physics of the used mass analyzer.

As the above-mentioned uncertainty is dependent on the *mz*, a fixed mass tolerance value for peak detection may result in peak merging and/or failure to detect peaks at certain *mz* range. An alternative way is to use a dynamic mass tolerance according to the uncertainty of acquired ions. Thus, the peak detection mass tolerance (*PDMT*), should be set as a function of the MF (usually defined in ppm unit) and MD. The peak detection mass tolerance in Da unit (PDMT_Da_) can be calculated by

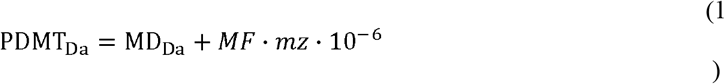

FTICR:

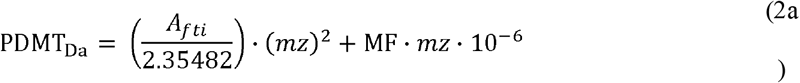

Orbitrap:

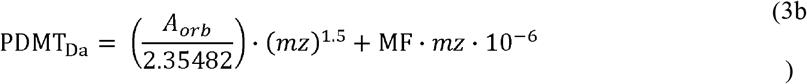

Q-TOF:

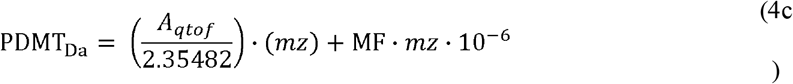

Quadrupole:

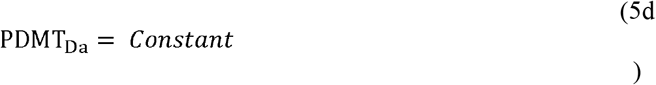

While the peak detection mass tolerance in ppm unit (PDMT_ppm_) can be calculated by

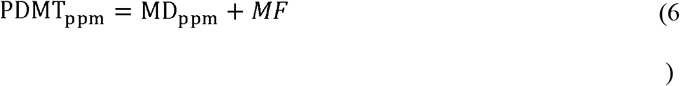

In what follows, we describe the equations for calculating *A*_*orb*_ in Orbitrap mass spectrometer. The equations for *A*_*qtof*_ in Quadrupole time-of-flight (*Q-TOF*) and *A*_*fti*_ in Fourier-transform ion cyclotron resonance (*FTICR*) mass spectrometers can be found in the Appendix.

### 2.2. Estimation of the mass dispersion according to the mass peak width

The variation of the *mz* defined by the MD is linked to the concept of the mass resolution or mass resolving power of a particular mass analyser. We first define the important concepts that are necessary to express the MD in terms of *mz* and mass resolution, based on the theory described in the book by *Hoffmann and Stroobant*,[31] it should be noted however that the terms may be defined differently in other sources.[32] Mass peak width (Δ*m*): full width at half maximum (*FWHM*) of mass spectral peak.

Mass resolving power (*R*): the observed mass (*m*) divided by the mass peak width (Δ*m*) at 50% height for a well-isolated single mass spectral peak, as illustrated in *Eq. (3)*.

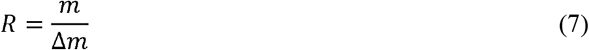

Therefore, the mass peak width (Δ*m*) changes according to mass resolving power (*R*) and the observed mass (*m*), as illustrated in *Eq. (4)*.

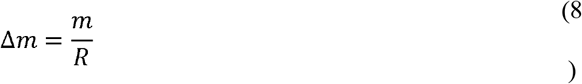

It follows from *Eq. (2)*, by replacing mass (*m*) with the mass-to-charge ratio (*mz*),

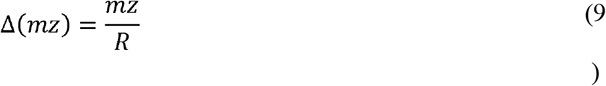

The underlying physical principle used in the estimation of *mz* is different for different types of mass analyzers. In an Orbitrap mass spectrometer, the frequency (*w*) is directly linked to the *mz* ratio, as illustrated in *Eq. (6)*

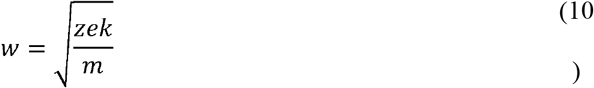

In which *e* represents the electron charge and *z* represents the number of charges of the ions. The letter *k* represents the field curvature, which is a constant value. Thus,

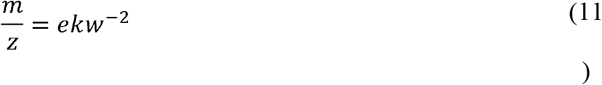

To find the relationship between the mass resolving power *R* and *mz*, we use the derivative of the mass with respect to the angular frequency from *Eq. (7)* and derive the *Eq. (8)* after rearrangement:

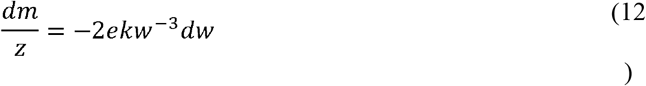

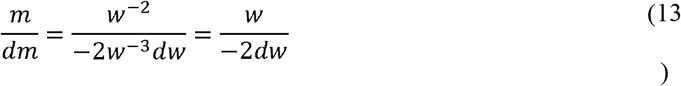

The ratio of mass to mass variation can then be obtained by dividing *Eq. (7)* by *Eq. (8)*

For a small variation, the derivative operator () and the difference used in numerical calculations (Δ) are interchangeable. The mass resolving power of the Orbitrap is further obtained by introducing the expression of the angular frequency from the *Eq. (6)* into *Eq. (9)* as follows

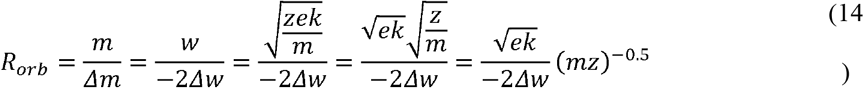

where Δm and Δw are 50% of the peak width measured on the mass scale and frequency scale, respectively. Several publications indicate the fact that the peak width in the frequency domain is almost a constant value: The seminal work of Makarov pointed to the fact that the errors of the frequencies are due to the construction of the instrument, rather than the experimental conditions.[33] The work of *Lössl et al*,[34] applied a fitting of the *Eq. (10)*, and showed an almost constant behaviour of the coefficient of *mz*^−0.5^. Thus, for fixed acquisition times, *Eq. (10)* indicates the mass resolving power R_orb_ is inversely proportional to the square root of *mz* ratio.[35] This can be expressed as:

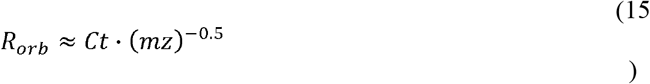

In which *Ct* indicates a constant value. Including *Eq. (11)* in *Eq. (5)*, the expression of the mass peak width for Orbitrap (i.e. Δ*mz*_*orb*_) becomes,

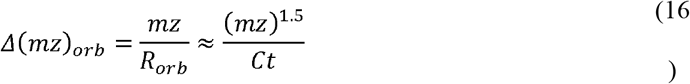

For a chosen reference *mz* (*mz*_*r*_),

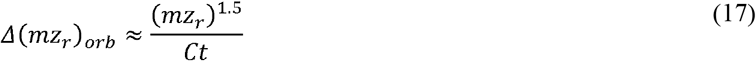

From *Eq. (12)* and *Eq. (13)* follows the equality,

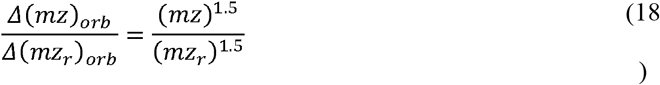

*Eq. (14)* can be further transformed into:

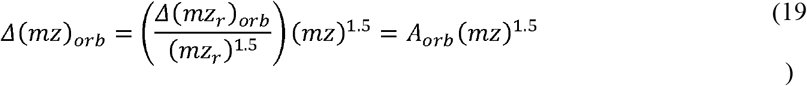

In *Eq. (15)*, *A*_*orb*_ is considered a constant value that can be calculated from the reference *mz*_*r*_ value and the reference resolving power *R*_*r*_, based on *Eq. (5)* becomes,

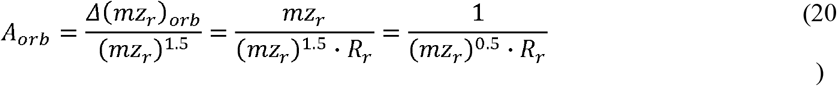

In which *R*_*r*_ indicates the reference resolving power at reference *mz*_*r*_. MD is usually quoted in terms of standard deviations (*σ*),[36] which can be estimated by the calculated mass peak width in *Eq. (15)*, according to the definition of a Gaussian-shaped function.[37]

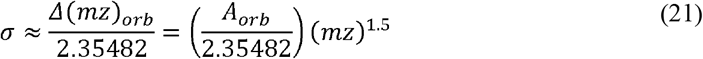

Finally, MD in Da unit (MD_da_) can be expressed as,

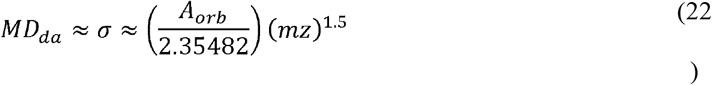

While the MD in ppm unit (MD_ppm_) can be expressed by

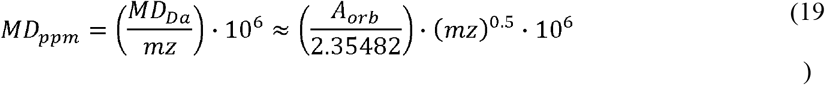

*Eq (18)* and (*19)* indicate that the mass dispersion changes as a function of mz, and consequently, according to the *Eq. (1)* and *Eq. (2)*, the PDMT should also change as a function of mz. The reference *mz* (*mz*_*r*_) is set to 200 Da, and the reference mass resolving power (*R*_*r*_) is set to 70000, according to *Eq. (16)*, *A*_*orb*_ equals 1.0102·10^−6^. The mass fluctuation (MF) is less than 1 ppm for a calibrated instrument at 200 Da.[30] We use MF = 1 ppm in the following equations. According to *Eq. (1)* and *Eq. (18)*

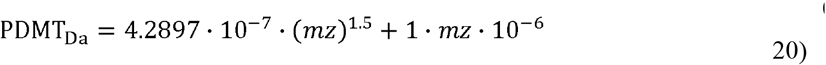

According to *Eq. (2)* and *Eq. (19)*

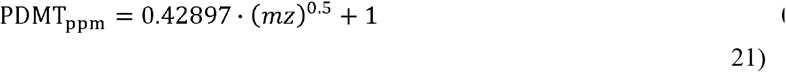

## 3. MATERIALS AND METHODS

### 3.1. Materials and Methods

#### Materials

LC-MS grade acetonitrile (ACN), methanol (MeOH), isopropanol (IPA) and chloroform were purchased from Biosolve BV (Valkenswaard, The Netherlands). Ammonium formate (AmF), formic acid (FA) and *tert*-Butyl methyl ether (MTBE) were purchased from Sigma Aldrich (St. Louis, MO). Various lipid standards were purchased from Avanti Polar Lipids, Inc. (Alabaster, AL). Heparin-anticoagulated plasma samples were obtained from adult patients at the University Medical Center Groningen (UMCG) in an anonymous manner and were combined to generate a standard plasma sample as described previously [38].

#### Deuterium lipid internal standard (IS) mixture preparation

20 different deuterium-labelled lipid IS and 4 deuterium-labelled lipid IS premix were selected to cover the major lipid classes and distributed evenly in *mz* and retention time range. All lipid standard stock solutions were diluted with chloroform: MeOH (1:1, v/v) and mixed to generate a lipid IS mixture with optimized concentrations for each standard to acquire adequate signal intensity (as listed in Table A.1). The lipid IS mixture was used to create a dilution series where concentration ratios were set to a factor of two starting from concentration 1 up to concentration 1/16.

#### Lipid extraction

Plasma lipid extraction was performed following the protocol of *Matyash et al*.[39] with slight modification. In brief, 60 μl of plasma was mixed with 300 μl of MeOH and sonicate for 10 min. Subsequently, 1000 μl MTBE was added and the mixture was kept under 25 °C on a shaker (900 rpm) for 30 min. Phase separation was induced by adding 190 μl ultrapure water. Then the mixture was centrifuged at 3000 RCF for 10 min and the 850 μl upper phase were transferred to a new tube. The re-extraction was performed by adding 600 μl MTBE/MeOH/ultrapure water (10:3:2.5, v/v/v) into the lower phase and 500 μl were collected after centrifugation to combine with the previous organic phase. The combined lipid extract solution was aliquoted into 6 tubes (190 μl per tube) to generate plasma lipid matrix. Different concentrations of lipid standard mixture were added to the plasma lipid extract aliquots and dried in a vacuum centrifuge under 45 °C. The dried lipid extracts were resuspended with 30 μl chloroform:MeOH:MQ (60:30:4.5, v/v/v) and further diluted with 90 μl (IPA:ACN:MQ 2:1:1 v/v/v) for LC-MS analysis.

#### LC-MS analyzis

LC-MS lipid analysis was performed on an Ultimate 3000 High-Performance UPLC coupled with a QExactive Orbitrap instrument (Thermo Fischer Scientific, Darmstadt, Germany). Chromatography separation was achieved with an Acquity UPLC CSH column [1.7 μm, 100□×□2.1 mm, (Waters Corporation, Milford, MA)] under 55°C with a flow rate of 0.4 ml/min. Mobile phase A was composed of ultrapure water/acetonitrile 40:60 (v/v), 10 mM ammonium formate and 0.1% formic acid. Mobile phase B contained ACN/IPA 10:90 (v/v) with 10 mM ammonium formate and 0.1% formic acid. The LC gradient was modified from *Damen et al*.[40] It started with 40% mobile phase B and raised to 43% mobile phase B in 2 min. The percentage of mobile phase B raised to 50% in the next 0.1 min and increased to 54% in next 9.9 min. Mobile phase B raised to 70% in 0.1 min and increased to 99% in 5.9 min and maintained at 99% for 1 min. Then the percentage of mobile phase B went back to 40% in 0.1 min and the system was equilibrated for 3.9 min before the next run started. The MS was set for positive mode and data-dependent acquisition. A full MS scan the range from 250-1750 Da was acquired at resolution 70,000 FWHM at 200 Da. AGC target was set to 1·10^6^, maximum injection time was 50 ms, MS1 scan was followed by up to 8 MS/MS events with a collision energy of 25 eV at resolution 17,500 FWHM at 200 Da. The precursor isolation window was set to 1.5 Da with the dynamic exclusion time of 6□s. The ionization settings were as follows: capillary voltage, +3.2□kV; capillary temperature: 320 °C; sheath gas/auxiliary gas: 60/20.

### 3.2. Computational Methods

#### Quality score calculation

Fig. A.4 shows the different concentrations of lipid IS mixture spiked in the plasma lipid extract aliquots. Thus, four fold changes are generated by comparing IS 1 with IS 1/16 (fold change 16), IS 1/8 (fold change 8), IS 1/4 (fold change 4), IS 1/2 (fold change 2). The t-statistics using standard two independent samples t-test between pairs of these four fold changes are calculated according to *Hoekman et al*.[29] Based on these t-statistics, the quality scores introduced in this paper were calculated. To get the distribution of these quality scores, in each of the concentrations, we randomly selected 9 out of the 10 replicates to calculate the t-statistics and the quality score. This selection was repeated for 20 times resulting in 20 quality scores for each comparison.

#### IS adducts confirmation

Each IS described in materials and method are annotated as 10 adducts types: M+H-H_2_O, M+H, M+NH_4_, M+Na, M+K, M+2Na-H, 2M+H, 2M+NH_4_, 2M+Na and 2M+K, the *mz* value were calculated accordingly. According to the calculated *mz* value, the EICs were visualized. Based on these visualizations, we confirmed the presence of 159 IS adducts. The criteria for confirmation of an EIC corresponding to an IS are: (1) the corresponding EIC should not show a bell-shaped peak signal in the blank samples; (2) the corresponding EIC should form a bell-shaped peak in both replicates of the IS mixture; (3) the apex of the bell-shaped signal (i.e. peak height) should pass the noise level of 10^5^ ion counts in the IS mixture and in the highest spiked-in concentration in human plasma samples; (4) the EIC should be identified in both pure IS mixture and IS mixture spiked plasma samples in at least in three spiked-in concentrations; (5) the EIC should follow the concentration fold changes in IS mixture spiked plasma samples in at least in three spiked-in concentrations. The EICs for the IS adducts are available in the Appendix. An example of EIC of a confirmed IS adducts can be found in Fig. A.5.

#### IS fragments confirmation

After a careful visual inspection of the collected LC-MS data, we also identified 416 IS fragments apart from the IS adducts described in the previous paragraph and included these in the IS list. The criteria for selecting these IS fragments are the same as the IS adducts. The EICs for the IS fragments are available in the Appendix. An example of EIC of an IS fragments included in the analysis is shown in Fig. A.6.

#### Lipidomics data processing

*Thermo Xcalibur^®^ software* (version 3.2.63, Thermo Scientific, Waltham, MA) was used for data acquisition (Fig. A.4). The acquired .raw thermo files were used directly by *Progenesis QI* (version 2.1) for peak picking, grouping and isotope filtration. The .raw data were further converted into .mzML format using *MSConvert* (Version: 3.0.18234) tool of ProteoWizard[41] package, the *Vendor Peak Picking* filter was selected to export *centroid* data. The .mzML files were sent to *mzMine* (version 2.40.1) for peak picking, grouping and isotope filtration. *XCMS* (version 3.8.2) also used .mzML files as input for peak picking and grouping for both the dynamic and fixed binning methods. The obtained results were further isotope filtered by *CAMERA* (version 1.43.2). The exported .csv files were used to assess the performances of the different LC-MS pre-processing tools. The fixed binning term reflects one parameter set in ppm used for linearly expanding EIC construction *mz* range in function of *mz*.

#### Dynamic binning implementation

In *XCMS*, the *centWave*[26] algorithm is one of the most commonly used peak detection algorithms, which uses a mass tolerance for EIC construction in *ppm* unit. In this study, the *centWave* algorithm in XCMS source code was modified to implement the dynamic binning approach, which is available in the Appendix. The *mz* of the dataset ranges from 250 Da to 1750 Da, thus the mass tolerance in modified XCMS source code was changed according to the presented theory.

## 4. RESULTS AND DISCUSSION

### 4.1. Model validation

To validate our model to set EIC construction threshold, we visually inspected the sampling interval in the profile mode dataset. The sampling intervals are the distances in *mz* between two adjacent acquisition points, which is determined by an algorithm implemented on the electronic card of Orbitrap instrument performing data processing with goal to have a constant number of sampling points for each peak in the mass spectrum. Since the number of sampling points in Orbitrap mass spectra is constant, then sampling interval increase should be in accordance with our theory on increasing peak width. *Fig. 2a* shows that the manually checked sampling intervals (red dots in the figure) increase proportional to 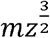. This increasing behavior of the sampling intervals set on the acquisition instrument and found in profile mass spectrum is following the peak width increase in function of *mz* described by *Eq. (15)*.

**Fig. 2.**
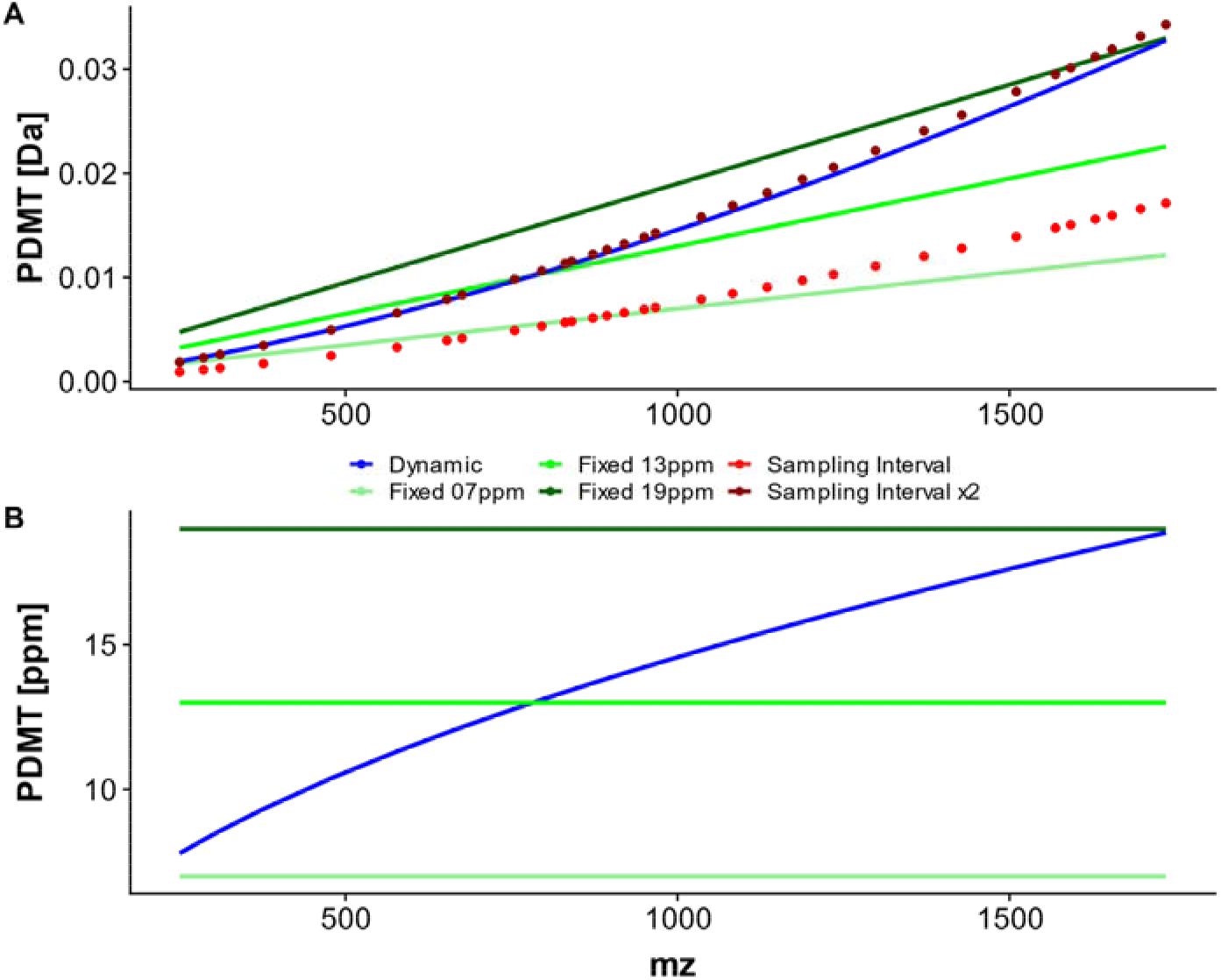
Validation of the theoretical model calculating PDMT in function of *mz* with sampling intervals of mass spectra in Orbitrap mass analyzer. (a) The manually checked sampling intervals (red dots in the figure) increase proportional with *mz* ^3/2^. This increasing behaviour of the sampling intervals is following the equation (15) of our theory to model peak width. (b) the PDMT *mz* tolerance in function of *mz* as implemented in dynamic binning method using *centWave* XCMS peak detection algorithm. The dynamic binning threshold is shown with blue line, while the fixed binning method originally implemented in XCMS with 7, 13 or 19 ppm threshold is shown as different green lines.

The exact number of sampling intervals defining the m/z interval which contains all ion intensity information for a peak in profile mode may vary depending on instrumental setting such as resolution. Empirically, this number can be estimated as 4. *Myers et al*[25] found, in centroided data, that most of the peaks in a mass spectra span typically between 0-3 sampling intervals. They chose a fixed *mz* tolerance value of 0.01 Da and 0.02 Da for Orbitrap and Q-TOF, respectively. This setting can ensure that most of the centroid maxima of the analysed isotopic ions in consecutive mass spectra (Fig. 1a) are included in the EIC construction. The *mz* tolerance value is estimated according to 1 sampling interval at the highest *mz*. In our model, the peak detection mass tolerance in Da unit (*PDMT_Da_*), calculated according to *Eq. (20)*, is between 1 and 2 sampling intervals (Fig. 2a). This dynamic setting for peak detection mass tolerance could, on one hand, avoid merging EICs at low *mz* and on the other hand avoid EICs splitting of one isotope peak at high *mz* values.

In the implemented dynamic binning method, the *mz* tolerance value in *centWave* was set to the dynamic *PDMT* value, while for the fixed binning method 7, 13 or 19 ppm (Fig. 2b) were used for peak detection for the entire *mz* range.

### 4.2. Comparison between the dynamic and fixed binning strategy in XCMS

Fig. 1 and 2 show examples when fixed binning method may fail to detect several ISs at higher *mz* ranges because limited ions are included for EIC construction and therefore the constructed EICs may not pass XCMS’s filtration parameters such as *firstBaselineCheck, prefilter* or *snthresh*. Fig. 3 shows two ISs that cannot be detected by using a fixed 7 ppm value of *mzTolerance* for EIC building, which was set following the guideline for setting XCMS parameters.[20] The lower part of the figure shows the scatter plot of *mz* and retention time, while the upper part shows the peak shape as a scatter plot of intensity and retention time. For example, the peak in Fig. 3a can be detected by using 13 ppm, 19 ppm with a fixed value of *mzTolerance* and by applying dynamic binning method. This peak should be detected in XCMS because 1) it matches M+Na adduct of IS 24:1(3)-14:1 CA, 2) it contains intensive signal in the *mz* range between 1676.236 and 1676.239 in Da, which 3) follows a bell-shaped curve with tailing in the retention time dimension with retention time range between 1099 and 1134 in seconds; 4) it has high intensity values, namely the area under the chromatographic peak is 1.648·10^8^, which becomes 1.646·10^8^ after baseline correction, and the maximum intensity measured in the *mz* and retention time rectangle containing the peak is 6.441·10^6^ (Table 1). Peaks in Fig. 3b shows similar properties as in Fig. 3a.

**Fig. 3.**
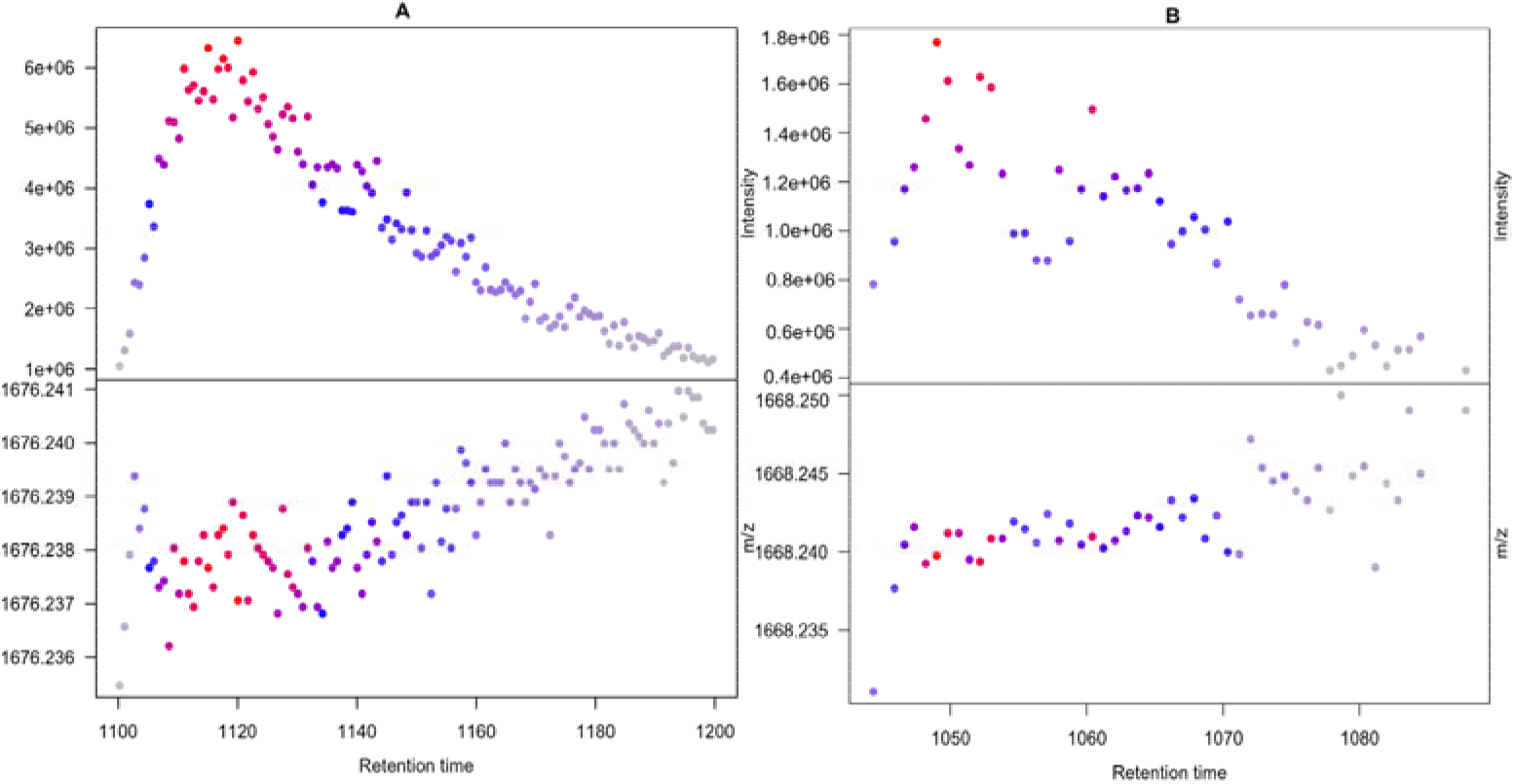
Examples of IS peaks not detected with the default 7 ppm fixed EIC construction threshold by XCMS. The lower part of the figure shows the scatter plot of *mz* and retention time, while the upper part shows the peak shape as a scatter plot of intensity and retention time.

**Table 1.** Examples of peak missing and improving quantification.

| method | $mz$ | $mzmin$ | $mzmax$ | $rt$ | $rtmin$ | $rtmax$ | $into$ | $intb$ | $maxo$ |
| --- | --- | --- | --- | --- | --- | --- | --- | --- | --- |
| Details of the missing peaks in Fig. 3 |  |  |  |  |  |  |  |  |  |
| Dynamic | 1676.238 | 1676.236 | 1676.239 | 1118 | 1099 | 1134 | $1.648 \cdot 10^8$ | $1.646 \cdot 10^8$ | $6.441 \cdot 10^6$ |
| Dynamic | 1668.240 | 1668.239 | 1668.242 | 1050 | 1045 | 1057 | $1.528 \cdot 10^7$ | $1.480 \cdot 10^7$ | $1.771 \cdot 10^6$ |
| Details of the peaks with improved quantification in Fig. 4 |  |  |  |  |  |  |  |  |  |
| Fixed 7ppm | 612.5013 | 612.5009 | 612.5042 | 843 | 838 | 846 | $1.241 \cdot 10^7$ | $1.241 \cdot 10^7$ | $3.498 \cdot 10^6$ |
| Dynamic | 612.5014 | 612.5009 | 612.5063 | 843 | 838 | 847 | $1.278 \cdot 10^7$ | $1.276 \cdot 10^7$ | $3.498 \cdot 10^6$ |
| Fixed 7ppm | 776.5780 | 776.5774 | 776.5785 | 415 | 404 | 423 | $5.628 \cdot 10^7$ | $5.613 \cdot 10^7$ | $7.296 \cdot 10^6$ |
| Fixed 7ppm | 776.5882 | 776.5860 | 776.5893 | 424 | 422 | 428 | $3.798 \cdot 10^6$ | $3.798 \cdot 10^6$ | $1.063 \cdot 10^6$ |
| Dynamic | 776.5783 | 776.5765 | 776.5883 | 415 | 404 | 425 | $5.758 \cdot 10^7$ | $5.716 \cdot 10^7$ | $7.296 \cdot 10^6$ |

$mz$ = mass to charge value; $mzmin$ , $mzmax$ = minimal and maximum $mz$ ; $rt$ = measured retention time in seconds; $rtmin$ , $rtmax$ = minimal and maximum $rt$ ; $into$ , $intb$ = intensity calculated based on area before and after baseline correction; $maxo$ = intensity calculated based on height;

Even though the constructed EICs pass XCMS’s filtration parameters, the quantification value of detected peaks may be less accurate compared to dynamic binning, because of less accurate EIC construction. The two peaks in Fig. 4 can be detected by using both fixed binning and dynamic binning method, but the quantification values are different (Table 1). For example, the highlighted point with black square in Fig. 4a is included only by the dynamic binning, while from the LC profile (upper part of Fig. 4a) it is obvious that the dots is part of the bell-shaped curve. As a result, the area of the peak detected with the dynamic binning method is higher than the one obtained with the fixed binning (i.e. 1.278·10^7^ compared with 1.241·10^7^). Fig. 4b shows an example of peak splitting using fixed binning peak detection. The two green rectangles indicate that the fixed binning method detected this peak twice at retention time 415 and 424 respectively; while dynamic binning method only detected correctly one peak at retention time 415, as indicated by the blue rectangle.

**Fig. 4.**
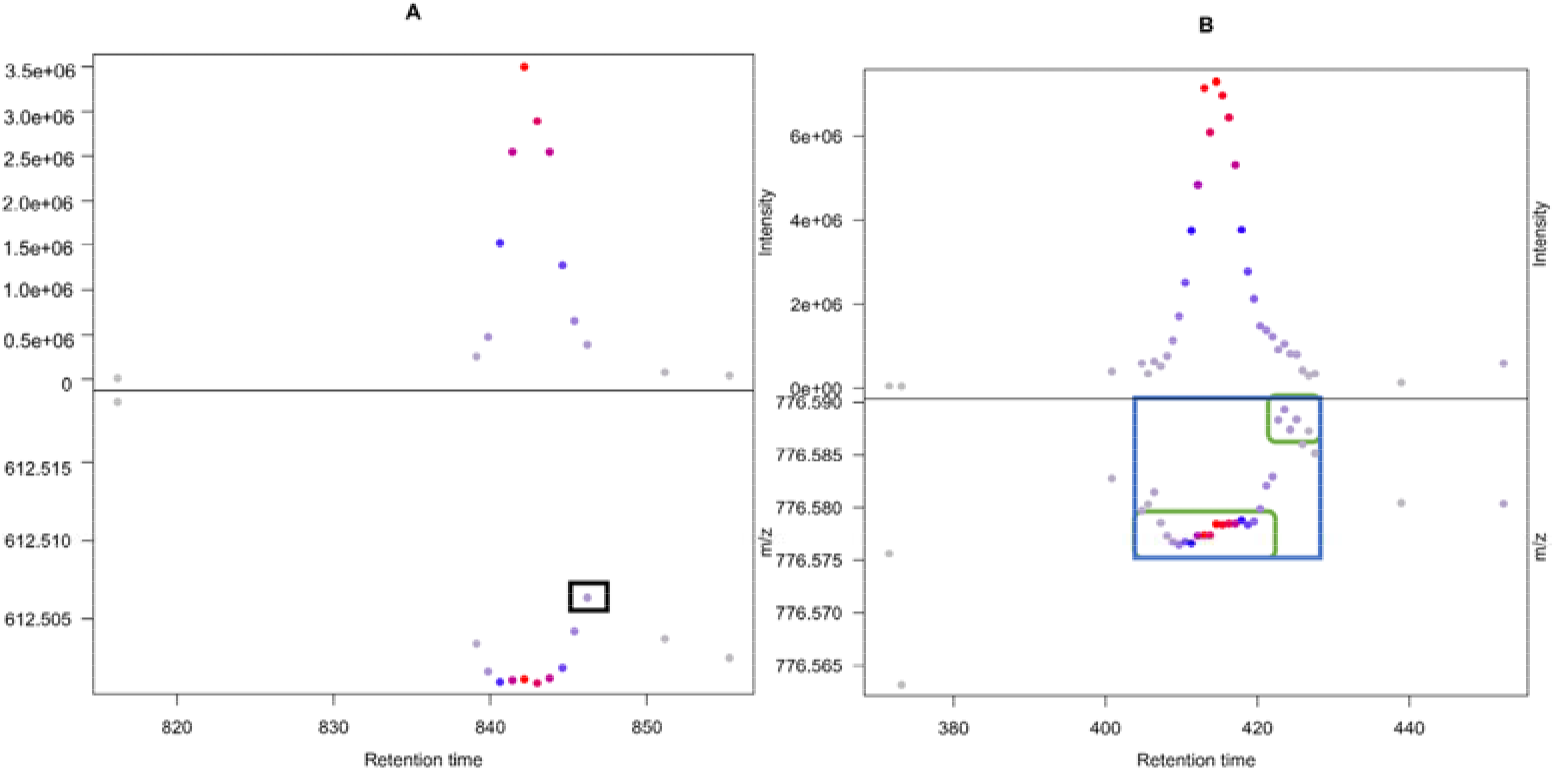
Examples of improved quantification by dynamic binning algorithm. Both peaks in the figure can be detected by using both fixed binning and dynamic binning method, but the quantification values are different. (a) The highlighted point is included only with the use of dynamic binning, while it is missing with fixed binning using 7 ppm threshold. This result in a higher area of a peak detected with the dynamic binning method compared to the one obtained with the fixed binning. (b) An example of peak splitting in fixed binning method with 7 ppm threshold. The two green rectangles indicate the two peaks detected by the fixed binning method, while dynamic binning method only detected correctly the one peak indicated by blue rectangle.

The difference between the number of detected peaks is small and from the presented data visualization of few examples is not obvious how dynamic binning performs at the whole dataset level. To have an overview of the difference of quantification between dynamic binning and fixed binning method considering all the peaks, we plotted the Bland–Atman plot of the quantification with dynamic and static EIC binning as shown in Fig. 5. The figure shows the comparison in concentration 1/16, 1/8, 1/4, 1/2, and 1 respectively. In each concentration, the x-axis shows the mean of the natural logarithm of peak intensities obtained with the dynamic and fixed binning method (7 ppm), and the y-axis represents the differences of the natural logarithms of the peak intensities obtained with both methods (i.e. log(*I*_*Dynamic*_)_*i*_ − log(*I*_*Fixed*_)_*i*_, for the i^th^ peak). In each concentration, the bias of the two methods (dashed blue line) is considerable showing larger quantification values for dynamic binning (i.e. blue dashed line above 0), which can be further confirmed by larger number of dots above the upper limit of agreement (dashed green line) compared to the lower limit of agreement (red line) both as expressed as two standard deviations from the overall center (dashed blue line). In order to assess if the higher quantification values lead to more accurate quantification, we have applied a quality score introduced by *Hoekman et al*.[29] that assess ability to identify differential spiked-in compounds in samples with the same complex lipid background.

**Fig. 5.**
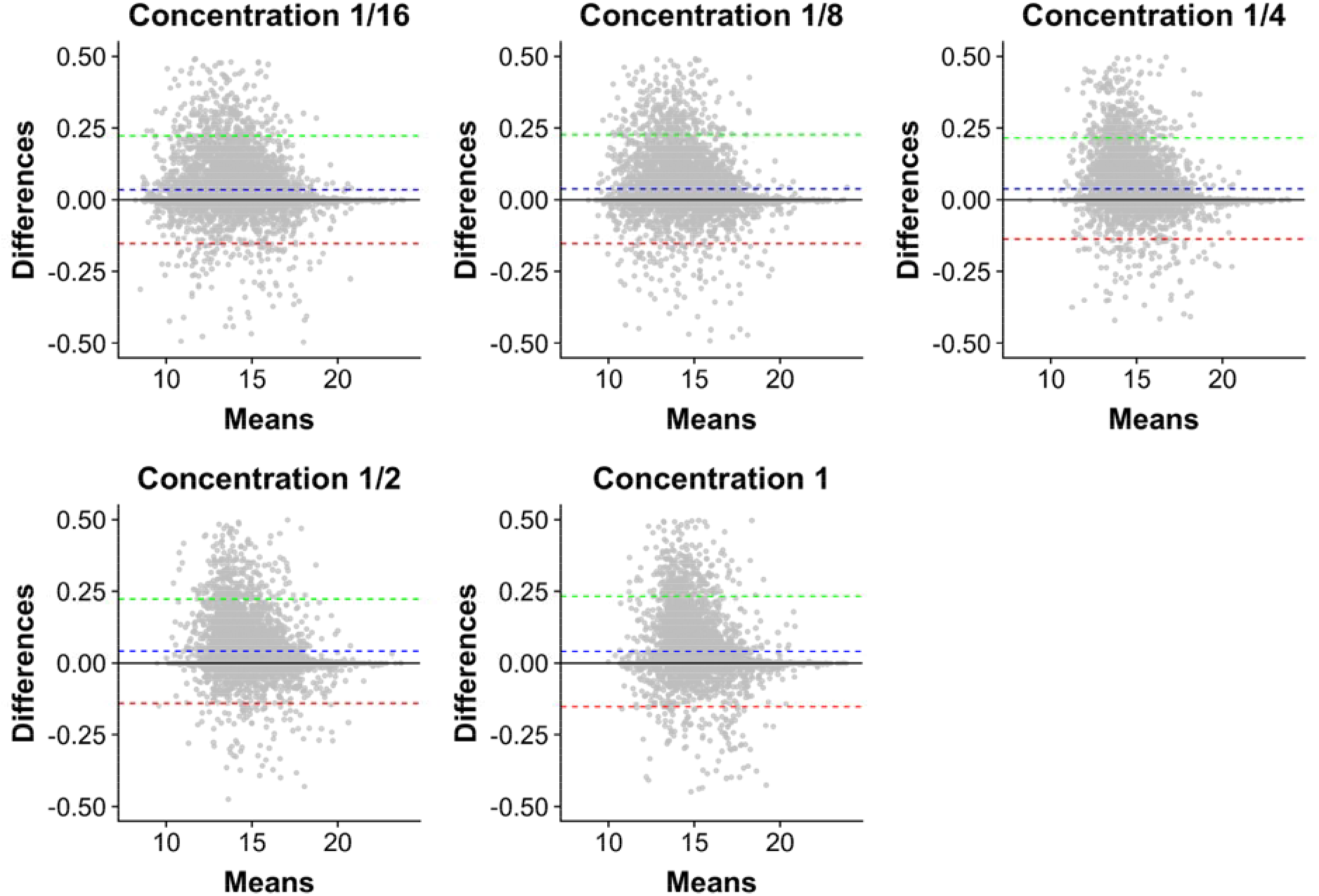
Bland-Altman plot showing the quantification differences between the dynamic and fixed bunning with 7 ppm EIC construction threshold for datasets with IS spiked with 1/16, 1/8, 1/4, 1/2, and 1 dilution level. In each plot, the x-axis shows the mean of the natural logarithm of peak intensities of obtained with the dynamic and fixed binning method, and the y-axis represents the differences of the natural logarithms of the peak intensities obtained with both methods (i.e. log(*I*_*Dynamic*_)_*i*_ − log(*I*_*Fixed*_)_*i*_, for the i^th^ peak). In each dilution level, the bias of the two methods (blue line) is above zero, which is further confirmed by larger number of dots above 0 on the y-axis. This indicates that quantification of the peaks obtained with dynamic binning are higher than quantification of the peaks obtained with fixed binning.

Fig. 6. shows the quality scores distribution of the studied pipelines for four different concentrations of fold changes (16, 8, 4 and 2 respectively). As indicated by the asterisks, in all these four different fold changes, the dynamic binning method achieved a quality score significantly higher than fixed binning method with 7 ppm (p-values 2.4·10^−4^, 1.9·10^−5^, 2.7·10^−4^ and 9.8·10^−3^ of two-samples Wilcoxon tests). The dynamic binning method improved XCMS’s ability to find the biomarker from the background was also compared to other fixed PDMT thresholds such as 13 and 19 ppm. The results show that dynamic binning overperformed original XCMS implementation.

**Fig. 6.**
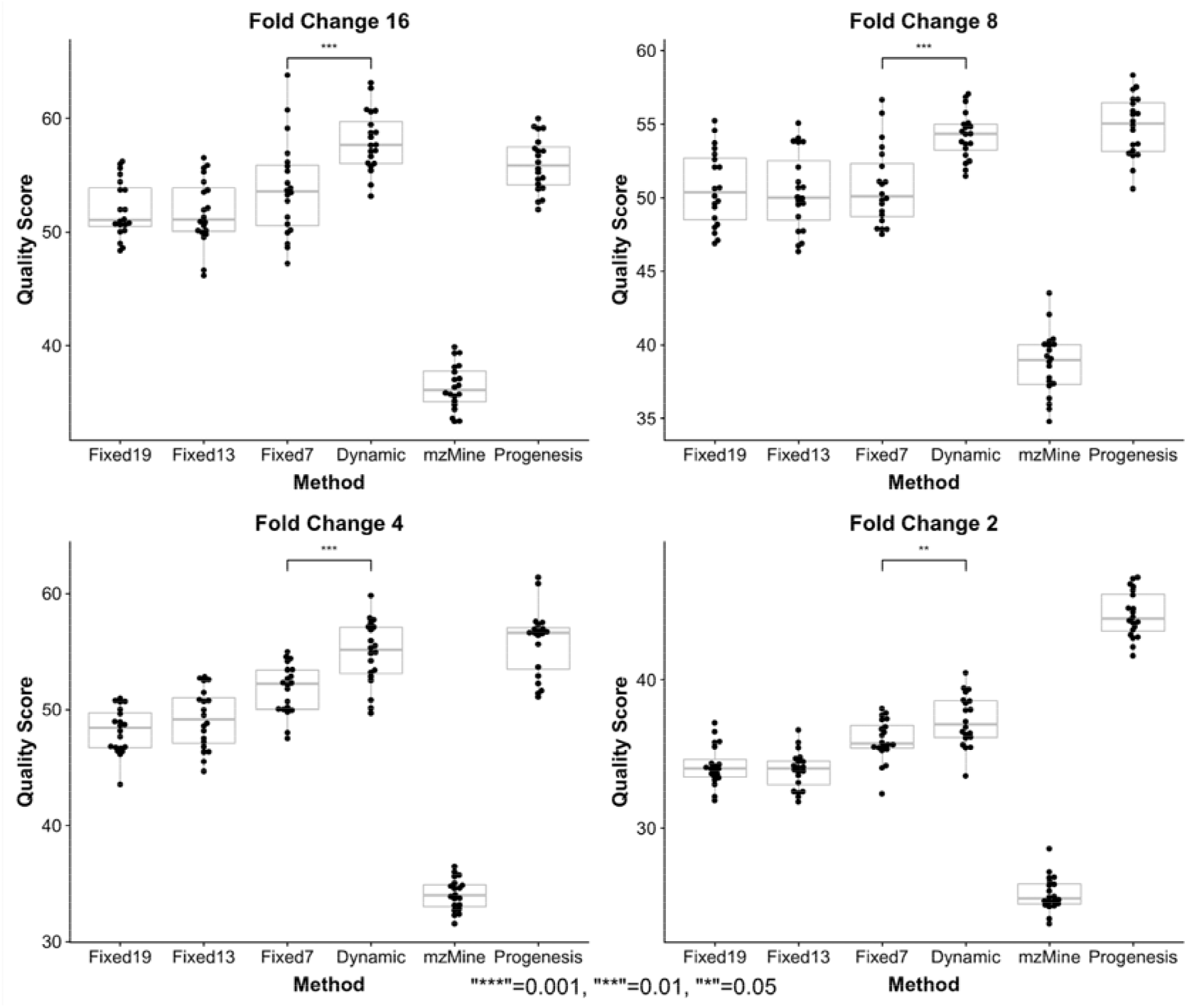
Box plot showing quality score, which measure the pipeline abilities’ to identify accurately differential peaks. The figure shows the quality scores distribution of the studied pipelines for four different fold changes (16, 8, 4 and 2 respectively), where dilution series of 1/16, 1/8, 1/4 and 1/2 are compared to dilution series of 1. The asterisks indicate the p-values (2.4·10^−4^, 1.9·10^−5^, 2.7·10^−4^ and 9.8·10^−3^) of two-samples Wilcoxon non-parametric tests performed between dynamic binning and fixed binning with threshold of 7 ppm. It is also obvious that dynamic binning method is performing better than fixed binning with 13 and 19 ppm thresholds and that the dynamic binning method improves XCMS’s quantification performance. Dynamic binning is performing much better than mzMine and perform similarly as Progenesis in 3 out 4 analysis.

The quality score assessing the performance to identify spiked-in features in stable molecular background is a cumulative score, i.e. it is also worth exploring the ranking of the spiked-compound-related features amongst the features identified to be differential between spiking levels. The cumulative quality scores in function of ranked list of discriminating features is shown in Fig. 7. In this figure, a binary heat map is included above the x-axis. Each row in this heat map indicates the differential features ranked from the most differential (t-statistics) to the weaker ones from left to right. In these, the colored bars indicate features related to the spiked-in compounds, which features are contributing to the quality cumulatively scores as indicated by the y-axis of the line plots. The white bars, on the contrary, indicate the features that correspond to compounds from the background sample which are non-discriminatory between the different spiked-in concentration levels. These non-discriminatory features will lower the increase in quality score for subsequent lower-ranked IS features. For example, the dynamic binning method contains more coloured bars than white bars amongst the most discriminant features at the left part of the plots, which indicates it detected more differentially spiked-in IS-related features as reflected in the quality score. This result indicates that dynamic binning method detect differential features more accurately as indicated by the higher number of spiked-in related features occurring amongst the most discriminative features compared with fixed binning. The two factors mentioned above, i.e. (1) larger number of coloured cells which (2) accumulate more abundantly at the left side of the plot, result in its higher cumulative score compared with fixed binning method at 7 ppm, 13 ppm and 19 ppm.

**Fig. 7.**
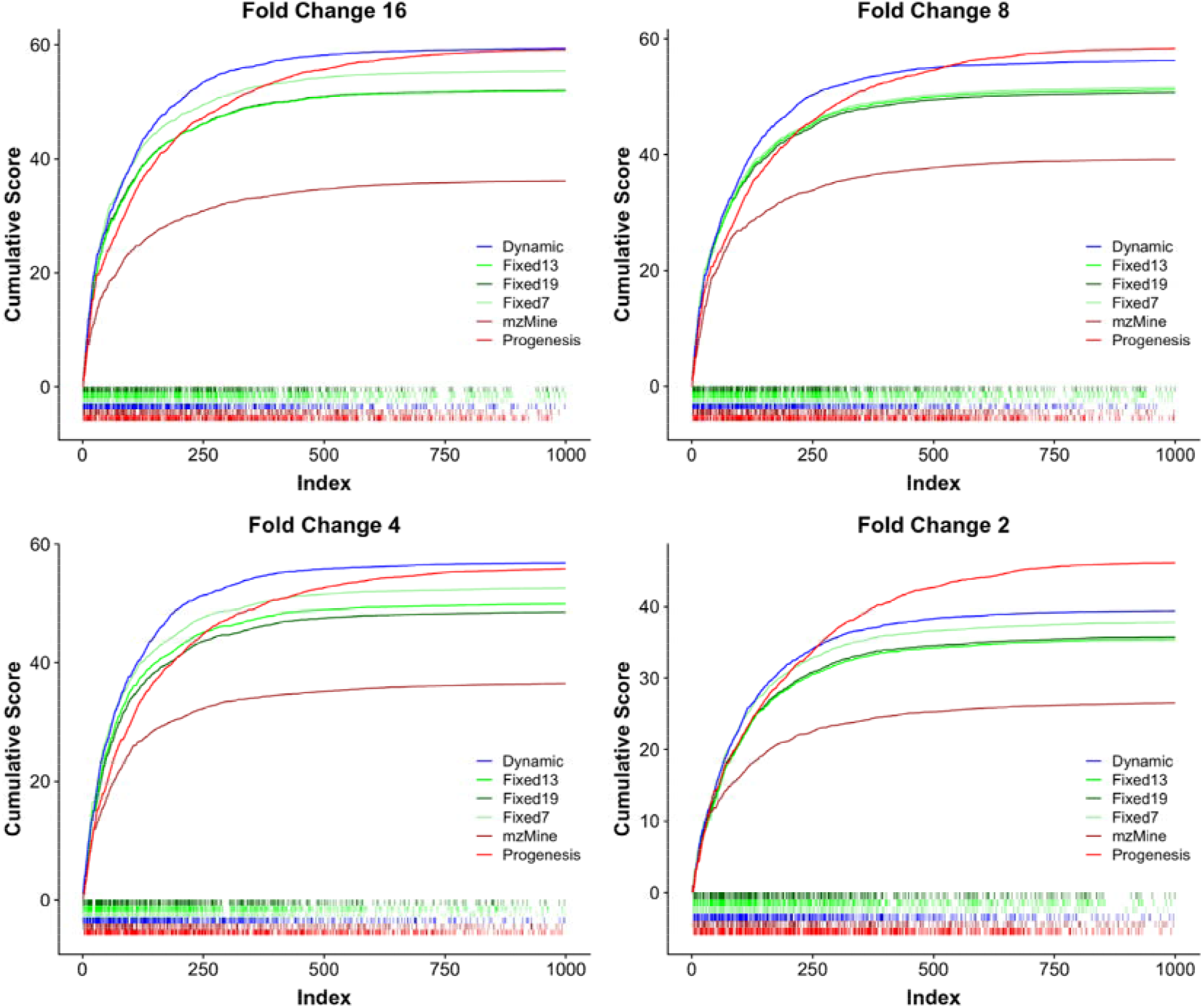
Comparison LC-MS pre-processing pipelines using plot of cumulative score development in function of ranked differential features. Differential metric of features is based on t-statistics calculated between two IS spiked-in levels and the theoretical fold change between spiking levels are indicted in the title of each plot. A binary heat map showing the ranked list of discriminating features from left (most discriminating) to right (less discriminating) is included above the x-axis of the cumulative score development plot. Each row in this heat map indicates the matched features related to the IS indicated as colored cells, which are increasing the cumulative quality score in the y-axis. The white cells indicate the features not related to IS and therefore these features 1) are not contributing to the cumulative quality score increase and 2) will lower the increase of the cumulative score for any subsequent higher rank IS related features. The dynamic binning method contains more colored cells than white ones amongst the most discriminant features in the left of the heat map, which indicates it detected more IS-related features amongst the most differential features.

### 4.3. Comparison between XCMS, mzMine and Progenesis

For a consistent comparison between XCMS, Mzmine and Progenesis, the quantitative feature tables containing matched features across all chromatograms in the dataset were used. The parameters for peak detection, grouping and isotope filtration can be found in Table A.2, A.3 and A.4. Fig. 6 shows that Progenesis and dynamic binning XCMS have constantly high-quality score values for comparison of spiked-in levels FC16, FC8, FC4 and FC2, followed by fixed binning XCMS and mzMine. The results indicate Progenesis and dynamic binning XCMS has a strong ability to find spiked-in compound related discriminating features and identify less background peaks having the same level across all spiked-in levels.

Apart from the aspect of finding more accurately features with different levels, it is also worth evaluating the precision of the quantification of different methods. Fig. 8 shows the scatter plot of log_2_ fold change (x-axis) versus log_10_ average abundance of a matched features (y-axis). The y-axis range of XCMS and mzMine is similar (5-11), while Progenesis is quite different (3-9), which indicates differences in the quantification metric. To assess if pipeline preserve the quantitative order of peaks, EICs (Fig. A.7) of three ISs were made. We labeled these three compounds as Compound A (17:0-17:1-17:0 D5 TG), Compound B (18:1-d7 MG) and Compound C (20:0-20:1-20:0 D5 TG) where the letter indicate the area abundance in decreasing order as shown in Table A.5. This table contains abundances calculated as area under the EIC curve from XCMS and mzMine, while in case of Progenesis, the “raw abundance” was used, which is the sum of all ion intensity in rectangle of all isotopes of a compound, where the rectangle are defined in an aggregated ion intensity map. The identified rectangles are then used in all aligned chromatograms to sum up the intensities and provide compounds quantitative values. The three compounds selected as example share the same abundance trend in XCMS and mzMine (i.e. 26.206·10^7^, 3.214·10^7^ and 1.980·10^7^ for compound A, B and C respectively in XCMS), while it is different in Progenesis (i.e. 12.425·10^7^, 0.089·10^7^ and 23.802·10^7^ for compound A, B and C respectively). Multiple possible reasons may exists to explain the abundance difference such as that XCMS and mzMine used the algorithm of *CAMERA* for isotope filtration, in which usually the highest isotope peaks are used for abundance calculation, while Progenesis used the average of isotope clusters for the abundance calculation. It should also be that the peak are of Compound A and B differs around 1 order of magnitude, which is accurately captured by the dynamic peak picking of XCMS and mzMine, while this difference is almost two order of magnitudes in case of Progenesis pre-processed data. Another explanation can be that the aggregate LC-MS map is constructed from aligned chromatogram of all fold changes. In this setup, the highest spiking level determine the area rectangle where ion intensity summation for all identified isotopes is performed and this rectangle is larger that should be for lower spiking levels features and may include therefore ions that do not belong to the spiking features leading to intensity order change.

**Fig. 8.**
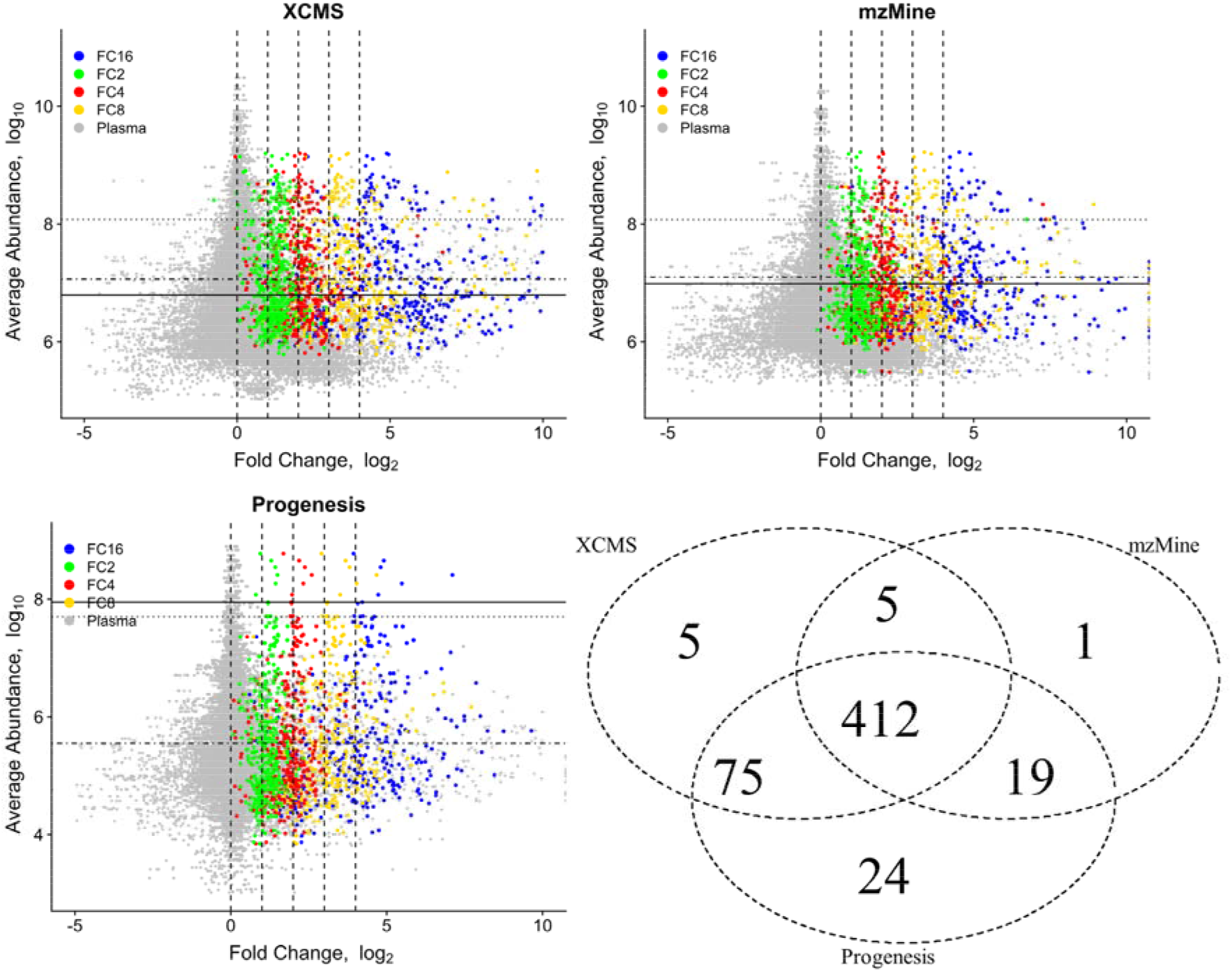
Ratio distributions between two spiking levels for each LC-MS pre-processing methods. The figure shows the scatter plot of fold change (x-axis, in log_2_) versus average abundance in the whole data set (y-axis, in log_10_). The y-axis range of XCMS and mzMine are similar (5-11), while Progenesis is different (3-9). In the x-axis, each IS is plotted four times with FC2, FC4, FC8 and FC16 labelled in green, red, yellow and blue respectively. The green, red, yellow and blue dots are located around log_2_ fold change of 1, 2, 3 and 4 respectively on the x-axis. The number of ISs detected by XCMS, mzMine and Progenesis is shown in the Venn diagram (bottom right plot), in which Progenesis uniquely detected 24 ISs, followed by XCMS (5 ISs) and mzMine (1 IS), which numbers are low compared to the number of features detected by all four methods (412). The features from LC-MS pipelines related to plasma are shown as grey dots, which fluctuate around log_2_ fold change of 0 in X-axis.

The x-axis of Fig. 8 reflects the fold change of detected features and each IS related features is plotted four times according to FC2, FC4, FC8 and FC16 and are labelled in green, red, yellow and blue respectively. In the plots the x-axis of the green, red, yellow and blue dots are located around the theoretical log_2_ fold change of 1, 2, 3 and 4 respectively. The number of ISs detected by XCMS, mzMine and Progenesis is shown in the Venn diagram, in which Progenesis uniquely detected 24 ISs, followed by XCMS (5 ISs) and mzMine (1 IS) are relatively low compared to the number of detected feature common to all pipelines (412 ISs). The features from XCMS, mzMine and Progenesis that are not related to ISs are labelled as feature that belong to the plasma (grey dots) are distributed around log_2_ fold change of 0 in x-axis. The fluctuation of the plasma related features in Progenesis is smaller compared with XCMS and mzMine. This is most probably due to the use of aggregate aligned map for feature detection, which perform relatively well in detection of compounds, which has low variability across samples, but may have difficulties for samples showing larger variability. In fact, the better performance of the dynamic binning peak picking is shown in Fig. 7 by the higher cumulative scores of this method over Progenesis for the most significantly different features i.e. features wish have the lowest rank and shown in the left part of the plot. Progenesis peak picking approach using the aggregate map allows detection of lower abundant peak due to the use of average signal across multiple LC-MS map for peak picking, which helps to identifying larger number of less differential peaks as indicated by the higher cumulative scores of Progenesis compared to dynamic binning XCMS approach taking into account all features for the score calculation.

To evaluate further the joint technical variance of LC-MS measurement and data pre-processing of three different methods, the distributions of the coefficient of variations (CV) were calculated based on the intensities of plasma related features. The full range of detected CV is shown as violin plot in Figure A.8. The density plot of CV between 0 and 1.5 (i.e. 150%) is shown in Fig. 9. These figures show that Progenesis has the most accurate feature quantification indicated by lower CVs and thus indicating lower variability of features in replicates of five individual concentration levels, while mzMine and XCMS have a wider distribution of CV. The better performance of Progensis is due to the use of aligned aggregate map for peak picking and feature detection, which can detect more accurately features with low variance as discussed previously.

**Fig. 9.**
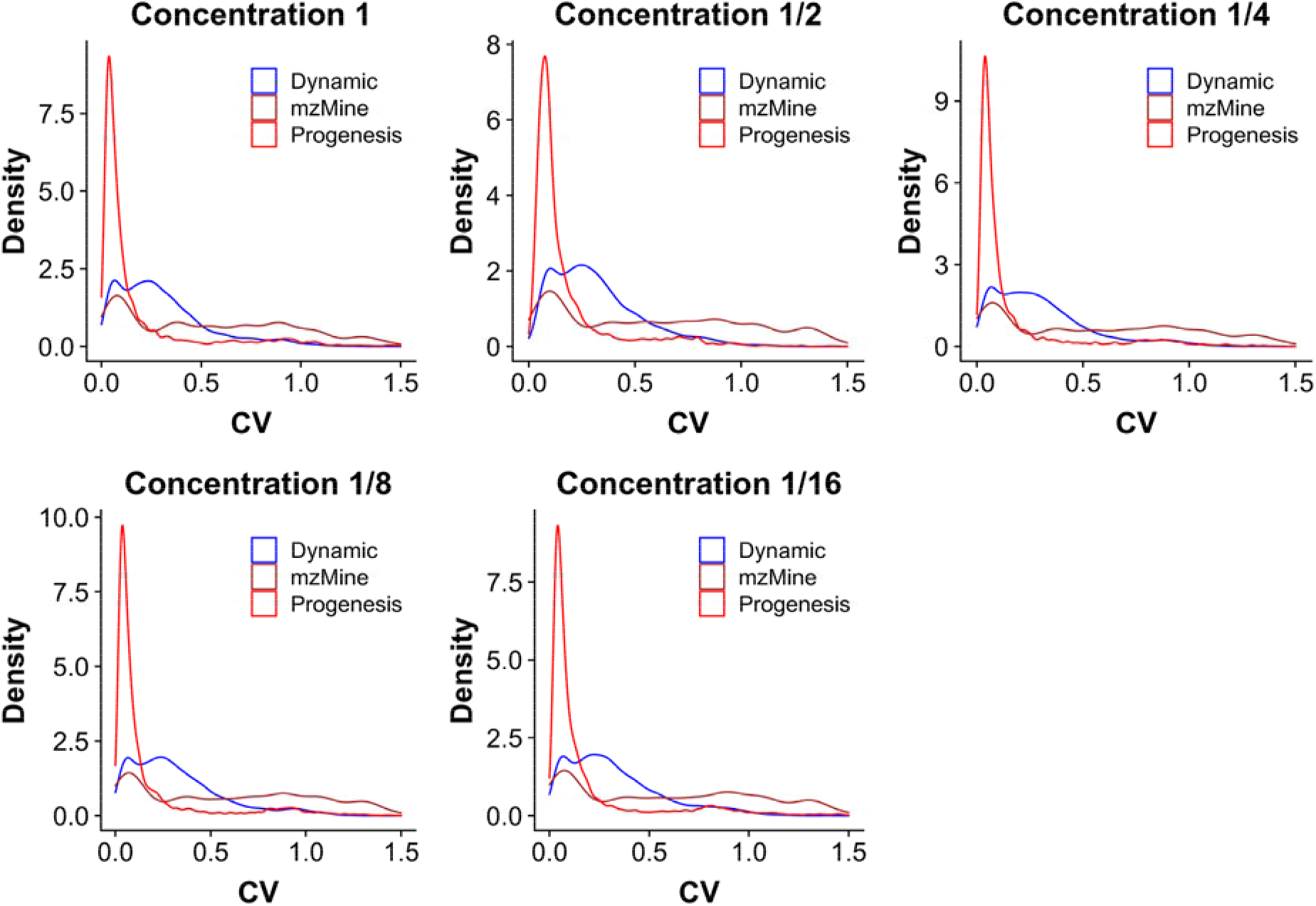
Density plot of coefficient of variation (CV) for 3 different LC-MS pre-processing methods using plasma related features. The plot shows the density of CV between 0 and 1.5 (150%). Progenesis shows the most precise feature quantification indicated by lower CVs in replicates of five individual concentration levels, while mzMine and XCMS have a wider distribution of CV.

## 5. CONCLUSION

Setting accurate PDMT is crucial for accurate peak detection. To this end, we suggested and implemented a dynamic method for more accurate setting of PDMT. Namely, the PDMT is proportional to *mz*^*2*^ for FTICR, to *mz*^*1.5*^ for Orbitrap, to *mz* for Q-TOF and a constant value for Quadrupole. This method improved XCMS performance by having fewer missing peaks (Fig. 3) and more accurate quantification (Fig. 4 and 5). As a result, the dynamic binning method shows a higher quality score (Fig. 6, maximum p-value: 9.8·10^−3^) and cumulative score (Fig. 7) compared with the fixed binning method. This indicates XCMS’s ability to find meaningful compounds (i.e. lipid biomarkers) is improved.

The results from the quality score and cumulative score also show Progenesis achieves similarly high quality scores and cumulative scores in all four fold changes to dynamic binning XCMS method, which is due to the use of aligned aggregate LC-MS for feature detection. Apart from the quality score and cumulative score, to evaluate the stability in quantification for XCMS, mzMine and Progenesis, we also determined the log_2_ fold change distribution (Fig. 8) and CV distribution (Fig. 9) based on the intensities of detected features. The results show that Progenesis has lower CVs for plasma related features, which indicates lower variability of features in replicates of five individual concentration levels, followed by XCMS and mzMine. The lower variability of Progenesis may be due to the unique quantification algorithm (Fig. A.7), however, this also altered the order of quantitative values of ISs (Table A.5). The use of aligned aggregate LC-MS maps for feature detection area may be beneficial to quantify peaks with low variability but can lead to artifacts in case of misaligned LC-MS maps and for features with larger variability. Dynamic binning peak picking in XCMS shows similar performance compared to Progenesis without the use of aligned aggregate maps and being exposed to the previous mentioned artifacts, we recommend it used in lipidomics and metabolomics quantitative profiling.

## ABBREVIATIONS

LC-MS(/MS): liquid chromatography-mass spectrometry (tandem mass spectrometry)
PDMT: peak detection mass tolerance
EIC: extracted ion chromatogram
MF: mass fluctuation
MD: mass dispersion
FWHM: full width at half maximum
Q-TOF: quadrupole time-of-flight
FTICR: Fourier-transform ion cyclotron resonance
IS: internal standard
Tr: tolerance range
CV: coefficient of variation

## ACKNOWLEDGEMENTS

The PhD research of Xiaodong Feng was supported by the China Scholarship Council grant No. 201708500094. This research was part of the Netherlands X-omics Initiative and partially funded by NWO, project 184.034.019.

## Appendices

- Appendix A. Tables Figures Equations Notes
- Appendix B. EIC visualization
- Appendix C. Improved XCMS source code

