## Appendix A for "Dynamic binning peak detection and assessment of various lipidomics liquid chromatography-mass spectrometry pre-processing platforms"

### 1. Equations

#### 1.1 Calculation of the mass peak width for (Q-)TOF mass analyzer

In (Q-)TOF mass spectrometer, mass-to-charge ratios are determined by measuring the time that ions take to move through a field-free region between the ion source and the detector. Indeed, before it leaves the ion source, an ion with a mass *m* and total charge of $q = ze$ is accelerated by a potential $V_s$. Its electric potential energy is then converted into kinetic energy. The observed mass resolution is derived from the relationship between ions' *mz* and flight time, as illustrated in *Eq. (1),* in which *e* represents the electron charge, *z* represents the number of charges of the ions, therefore the total charge of the ions *q* equals *ze*. $V_s$ represents the accelerating potential, *t* represents the time needed for the ion to travel the distance *L*.

$$\frac{m}{z} = \frac{2eV_s t^2}{L^2} \tag{1}$$

The *Eq. (2)* can be obtained by derivation of *Eq. (1)* based on time

$$\frac{dm}{z} = \frac{2eV_s 2tdt}{L^2} \tag{2}$$

The *Eq. (3)* can be obtained by dividing *Eq. (1)* by *Eq. (2)*

$$\frac{m}{dm} = \frac{t}{2dt} \tag{3}$$

Therefore, in reflectron mode, the resolution in (Q-)TOF mass spectrometer is equal to:

$$R_{qtof} = \frac{m}{\Delta m} = \frac{t}{2\Delta t} \approx \frac{L}{2\Delta x} \tag{4}$$

where ($\Delta m$) and ($\Delta t$) are the peak width measured at the 50% level on the mass and time unite, respectively. $\Delta$x is the thickness of an ion packet approaching the detector. Therefore, the (Q-)TOF mass spectrometer has a mass resolving power $R_{qtof}$ that remains constant when the mass increases. Here we use $C_{qtof}$ to represent the constant value in *Eq. (4).*

$$R_{qtof} = C_{qtof} \tag{5}$$

According to *Eq. (3)* in the main paper,

$$\Delta(mz)_{qtof} = \frac{mz}{R_{qtof}} = \frac{mz}{C_{qtof}} \tag{6}$$

The reference mass peak width can be calculated by:

$$\Delta(mz_r)_{qtof} = \frac{mz_r}{C_{qtof}} \tag{7}$$

Where the $mz_r$ represents the reference mass charge ratio. Thus,

$$\frac{\Delta(mz)_{qtof}}{\Delta(mz_r)_{qtof}} = \frac{mz}{mz_r} \tag{8}$$

Thus,

$$\Delta(mz)_{qtof} = \frac{\Delta(mz_r)_{qtof}}{mz_r} \cdot mz = A_{qtof} \cdot mz \tag{9}$$

where $A_{qtof}$ is a constant value reflecting quantities related to the reference *mz* that can be calculated by:

$$A_{qtof} = \frac{\frac{mz_r}{C_{qtof}}}{mz_r} = 1/C_{qtof} \tag{10}$$

#### 1.2 Calculation of the mass peak width for FTICR mass analyzer

In FTICR mass spectrometer, the angular velocity (*w*) is equal to

$$w = \frac{ze}{m}B \tag{11}$$

where *e* represents the electron charge, *z* represents the number of charges of the ions, *m* represents the mass of the ion, *B* represents the magnetic field, which is a constant value. As a result of this equation, the frequency and the angular velocity (*w*) depends on the *mz* ratio. Thus,

$$\frac{m}{z} = eBw^{-1} \tag{12}$$

The *Eq. (13)* can be obtained by derivation of *Eq. (12)* with *w*

$$\frac{dm}{z} = -eBw^{-2}dw \tag{13}$$

*Eq. (14)* can be obtained by dividing *Eq. (12)* with *Eq. (13)*

$$\frac{m}{dm} = \frac{w^{-1}}{-w^{-2}dw} = \frac{w}{-dw} \tag{14}$$

Therefore, the resolution in FTICR mass spectrometer is equal to:

$$R_{fti} = \frac{m}{\Delta m} = \frac{w}{-\Delta w} = \frac{\frac{ze}{m}B}{-\Delta w} = \frac{-eB}{\Delta w} \cdot (mz)^{-1} \tag{15}$$

where $\Delta m$ and $\Delta w$ are the peak width measured at the 50% level on the mass scale and frequency scale, respectively. The work of *Lössl et al.*[1] showed an almost constant behavior of the coefficient of $(mz)^{-1}$. Thus, for fixed acquisition times, *FTICR's* mass resolving power $R_{fti}$ is inversely proportional to the value of *mz*. Therefore, $R_{fti}$ can be estimated as:

$$R_{fti} \approx C_{fti} \cdot (mz)^{-1} \tag{16}$$

in which $C_{fti}$ is a constant value in FTICR. According to *Eq. (3)* in the main paper.

$$\Delta(mz)_{fti} = \frac{(mz)}{R_{fti}} \approx \frac{(mz)^2}{C_{fti}} \tag{17}$$

The reference mass peak width can be calculated by:

$$\Delta(mz_r)_{\mathrm{fti}} \approx \frac{(mz_r)^2}{\mathrm{C_{fti}}} \tag{18}$$

Thus,

$$\frac{\Delta(mz)_{fti}}{\Delta(mz_r)_{fti}} = \frac{(mz)^2}{(mz_r)^2} \tag{19}$$

Thus,

$$\Delta(mz)_{fti} = \frac{\Delta(mz_r)_{fti}}{(mz_r)^2} \cdot (mz)^2 = A_{fti} \cdot (mz)^2 \tag{20}$$

Where $A_{fti}$ is a constant value that can be calculated by:

$$A_{fti} = \frac{mz_r}{R_r \cdot (mz_r)^2} = \frac{1}{R_r \cdot mz_r} \tag{21}$$

In which $mz_r$ and $R_r$ indicate the reference *mz* and reference resolving power accordingly.

#### 2. Supplementary Figures

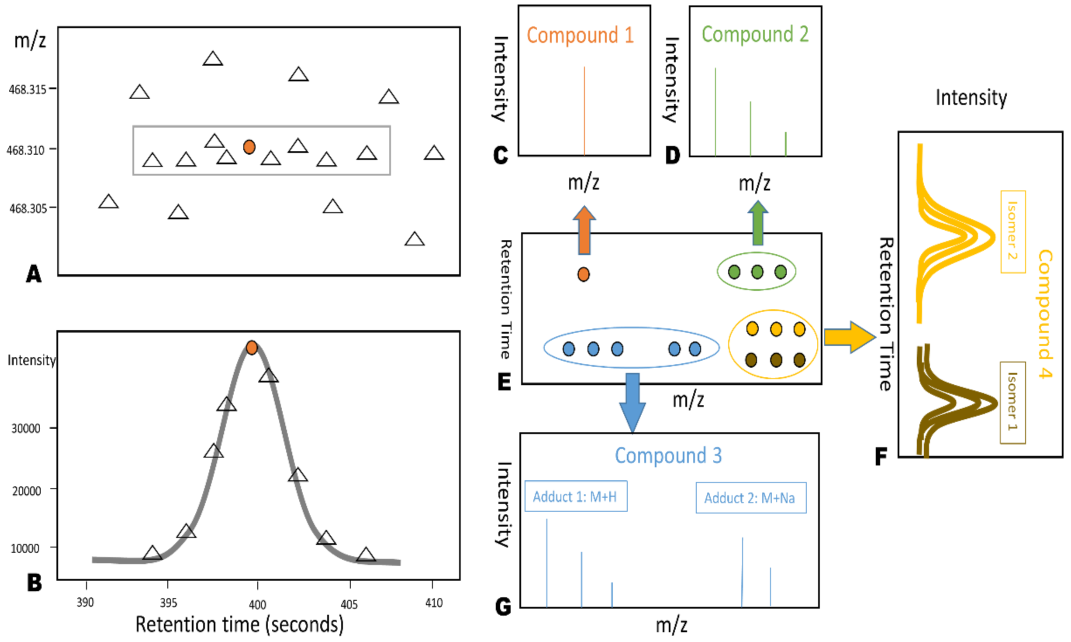

**Fig. A.1.** Scheme showing main aspects of peak detection. (**a**) shows an example of the construction of an EIC around an isotope peak in LC-MS map. (**b**) presents example of a bell-curve shape peak in the EIC. (**c**) shows an isotope (orange peaks) of compound 1, while (**d**) shows the isotope pattern of compound 2, which is composed of three isotopic peaks. (**e**) shows the ions of detected peaks in a piece of LC-MS map. (**f**) presents two isobaric structural isomers of compound 4 separated by liquid chromatography. Each isomer also shows an isotope pattern, which is the same in case of the same atomic composition of the molecules. (**g**) shows two adduct types of compound 3: $[M+H]^+$ and $[M + Na]^+$ with a distinct representative isotope pattern.

Peak detection is one of the most important steps in quantitative pre-processing of metabolomics LC-MS(/MS) workflows. Therefore, Fig. A.1 shows the general peak detection process. First, we define the following terms according to IUPAC nomenclature.[2]

**Extracted ion chromatograms (EICs)** are generated by selecting an *mz* value with an associated tolerance around and a retention time. (Note: for a clearer notation, we use *mz* across the entire article to represent *m/z*). The ions in the rectangle of Fig. A.1a show an example of the construction of an EIC. If the constructed EIC contains one or several bell-shaped curves as shown in Fig. A.1b, then these bell-shaped curves are identified as **peaks**. Peak detection is a process of selecting a point in retention time and *mz* which represent the location of a peak. For example, it is possible to choose the ion with the highest intensity (round marker in Fig. A.1b) to represent the location of a peak.

The *mz* coordinate of the peak can be used to determine the identity of the compound based on accurate mass. As shown in Fig. A.1c, the orange peak corresponds to one isotope of compound 1**.** An **isotope pattern** is defined as a set of isotope peaks related to one charge state and one adduct form of a compound. Isotopes of one compound have the same chemical formula (i.e. atomics composition) but have different isotopic composition and their relative abundance is determined by the natural abundance of stable isotopes such as $^{12}C$ and $^{13}C$. Fig. A.1d shows the isotope pattern of compound 2, which is composed of three isotopic peaks i.e. they are isotopologues. The identification of the isotope peak cluster is called feature detection or isotope filtration and is a common step applied before compound identification. After this step, only one peak is selected to represent a one charge state and adduct form of a compound, which is typically the isotope with the lowest *mz* called monoisotopic peak and is used to determine the identity of the compound. In this work, these retained peaks are referred to as **features.**

One compound may have multiple **features** corresponding to a various charge state and **adduct ions** but in small molecule analysis such as lipidomics and metabolomics charge state of ±1**,** which are defined as ions formed by the non-covalent interaction of a precursor ion with one or more atoms or molecules to form an aggregate ion. This aggregate ion contains all the atoms of the precursor ion and the additional atoms from the associated atoms or molecules (e.g. a Na+ adduct of a molecule (M) is represented as $[M+Na]^+$). Fig. A.1g shows that compound 3 contains two adduct types: $[M+H]^+$ and $[M+Na]^+$, and each adduct ion has a distinct representative **isotope pattern**. By using the *mz* of the charged precursor ion and the *mz* of the different adduct types (and the charge state), the compounds′ neutral masses can be calculated and be used for adduct types annotation. A compound may have structural isomers or diastereomers, which have different molecular 3D shape and physicochemical properties and may elute at different retention times.[3] As example

Figure A.1f shows two structural isomers of compound 4 separated by liquid chromatography.

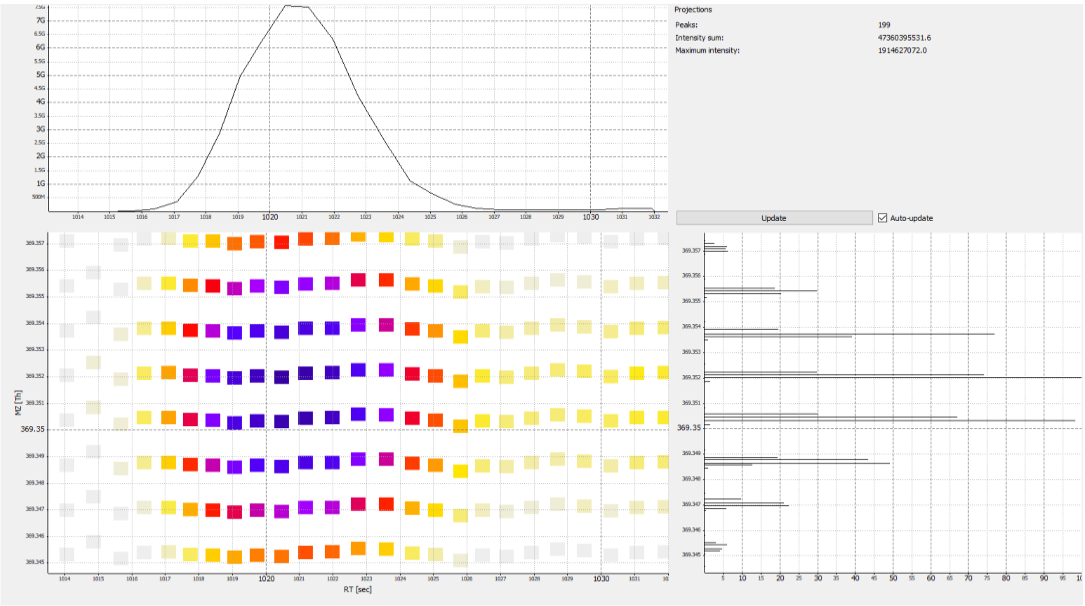

**Fig. A.2.** Mass fluctuation (MF) at low *mz*. The EIC (top left) around the *mz* 369.352 and the top view (bottom left) of the mass traces included in the selected EIC. Visualization made by TOPView tool in OpenMS.

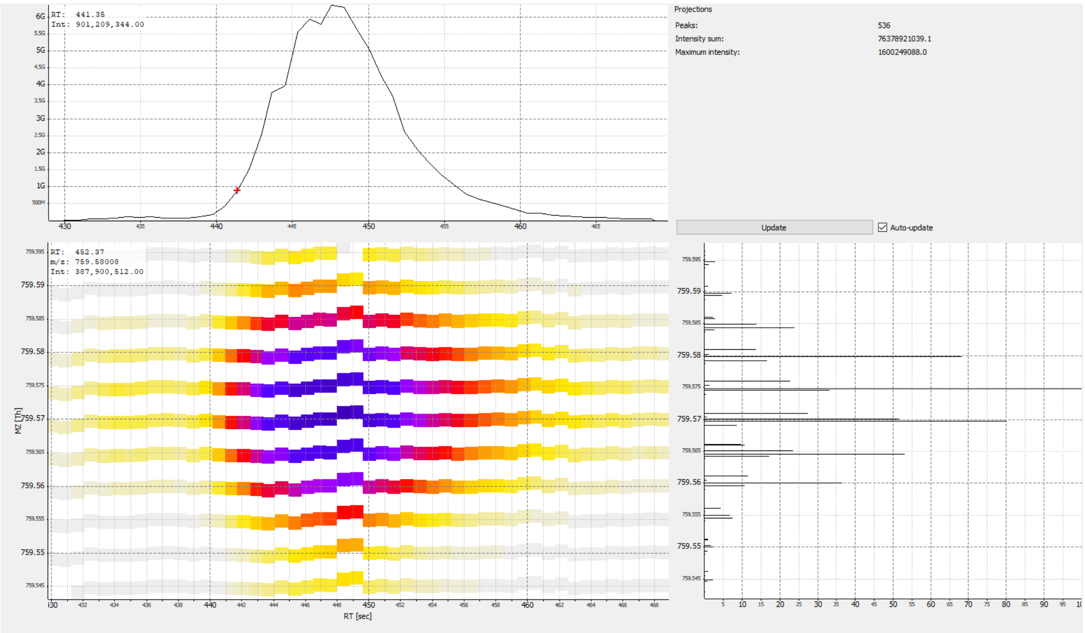

**Fig A.3.** Mass fluctuation (MF) at high *mz*. The EIC (top left) around the *mz* 759.570 and the top view (bottom left) of the mass traces included in the selected EIC. Visualization made by TOPView tool in OpenMS.

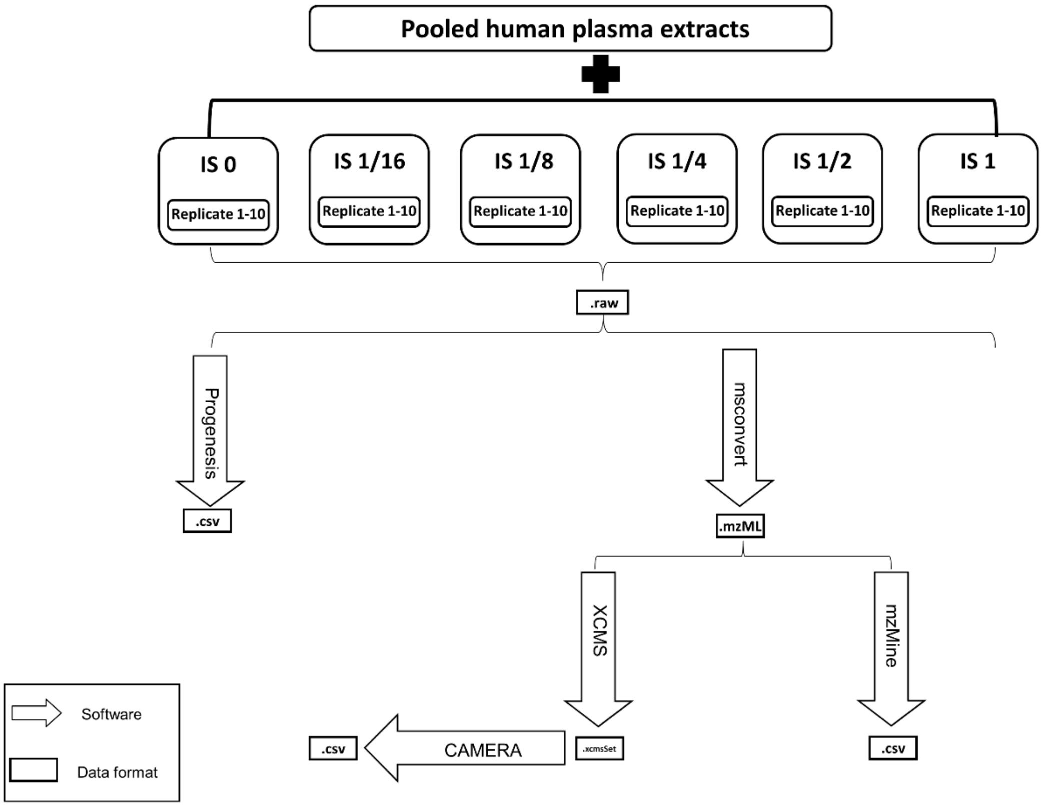

**Fig A.4.** Scheme of sample preparation and bioinformatics experimental design. Different concentrations of lipid internal standard mixture were added to the aliquots of one human plasma lipid sample. Four-fold changes between spiked-in concentration are generated by comparing IS 1 with IS 1/16 (fold change 16), IS 1/8 (fold change 8), IS 1/4 (fold change 4), IS 1/2 (fold change 2). The t-statistics using standard two independent samples t-test between pairs of these four-fold changes are calculated. Based on these t-statistics cumulative quality scores used to compare LC-MS pipeline to find differential peaks are calculated.

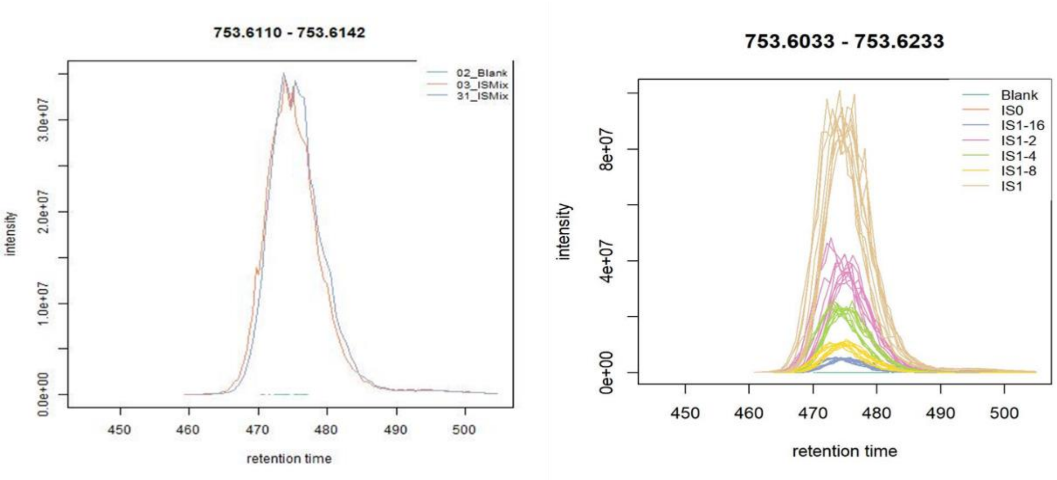

**Fig A.5.** Example of peak related to spiked-in internal standard. The criteria for confirmation of an IS related EIC are: (1) the corresponding EIC should not show a bell-shaped peak signal in the blank samples; (2) the corresponding EIC should form a bell-shaped peak in both replicates of the IS mixture; (3) the apex of the bell-shaped signal (i.e. peak height) should pass the noise level of $10^5$ ion counts; (4) the EIC should be identified in both pure IS mixture and IS mixture spiked-in human plasma aliquots ; (5) the EIC should follow the concentration trend in IS mixture spiked-in at different levels in human plasma aliquots.

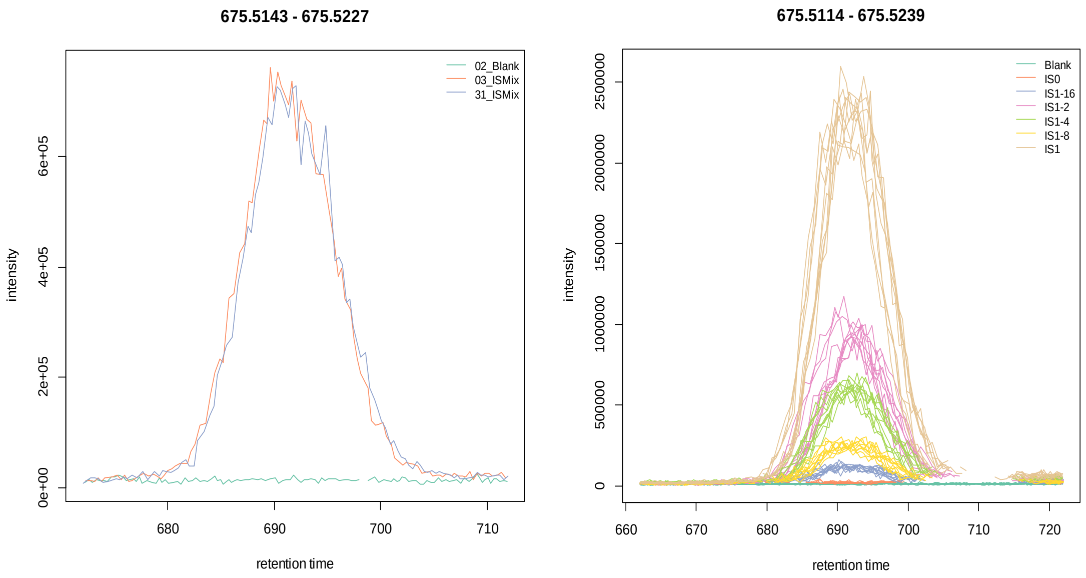

**Fig A.6.** Example of internal standard fragments. Left plot: EICs made for pure internal standard and a blank samples. Right plot: EICs related to internal standard spiked-in a human plasma aliquots. The internal standard fragment should be identified in both pure IS mixture and IS mixture spiked-in human plasma aliquots.

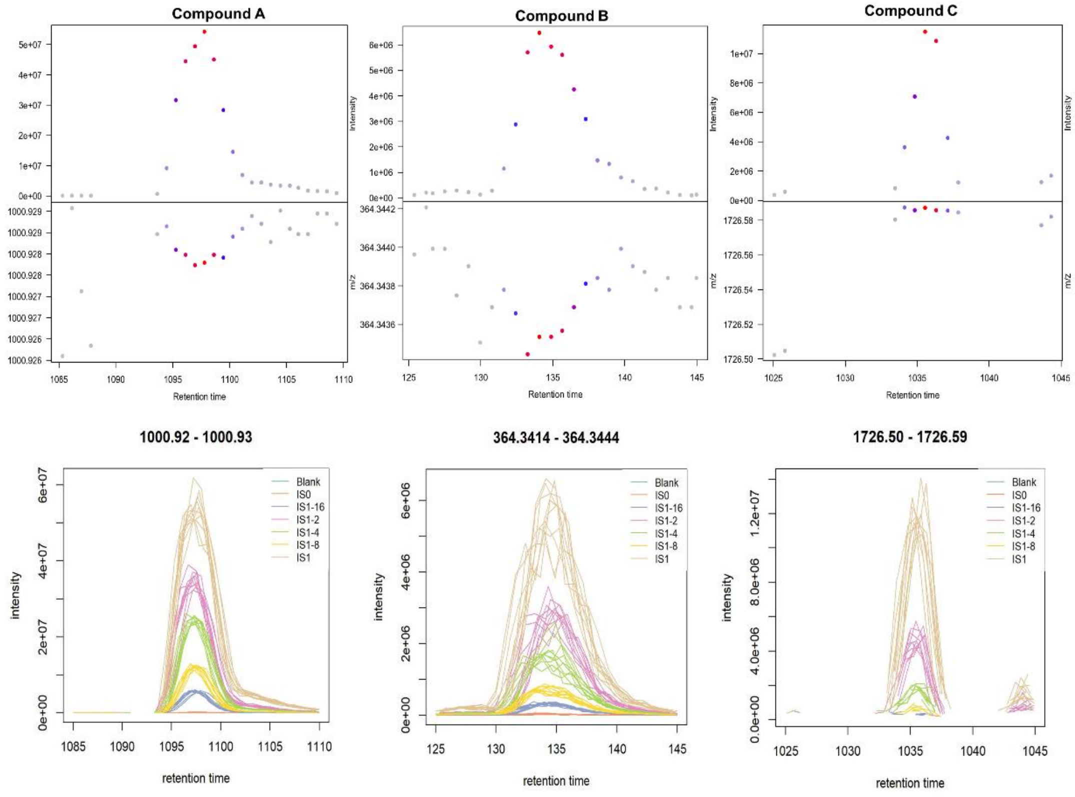

**Fig A.7.** EICs of three ISs shown in Figure 8. Compound A (17:0-17:1-17:0 D5 TG), Compound B (18:1-d7 MG) and Compound C (20:0-20:1-20:0 D5 TG) follow the abundance expressed as area under EIC curve in decreasing order.

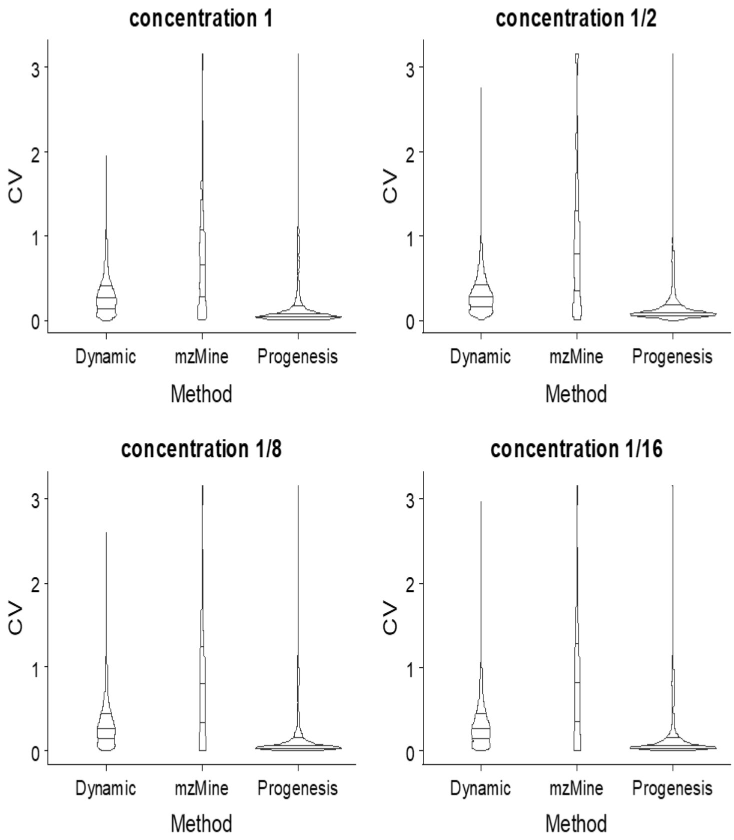

**Fig A.8.** Violin plot of the coefficients of variation of all the features for 5 IS spiked-in levels in human plasma aliquots.

### 3. Tables

**Table A.1.** 20 different deuterium-labelled lipid IS and 4 deuterium-labelled lipid IS premix were selected to cover the major lipid classes and distributed evenly in *mz* and retention time range. All lipid standard stock solutions were diluted with chloroform: MeOH (1:1, v/v) and mixed to generate a lipid IS mixture with optimized concentrations for each standard to acquire adequate signal intensity.

| **lipid Standard ID** | Standard name | Standard final concentration (pmol/ul) | | | | | |
|---|---|---|---|---|---|---|---|
| | | Conc 1 | **Conc 1/2** | **Conc 1/4** | **Conc 1/8** | **Conc 1/16** | **Conc 0** |
| **1** | 15:0-18:1-d7-PC | 2.5 | 1.25 | 0.63 | 0.31 | 0.16 | 0 |
| **2** | 18:1-d7 Lyso PC | 2.5 | 1.25 | 0.63 | 0.31 | 0.16 | 0 |
| **3** | C18(plasm)-18:1(d9) PC | 5 | 2.5 | 1.25 | 0.63 | 0.31 | 0 |
| **4** | 15:0-18:1-d7-PI | 7.5 | 3.75 | 1.88 | 0.94 | 0.47 | 0 |
| **5** | 15:0-18:1-d7-PE | 5 | 2.5 | 1.25 | 0.63 | 0.31 | 0 |
| **6** | 18:1-d7 Lyso PE | 10 | 5 | 2.5 | 1.25 | 0.63 | 0 |
| **7** | C18(plasm)-18:1(d9) PE | 5 | 2.5 | 1.25 | 0.63 | 0.31 | 0 |
| **8** | 15:0-18:1-d7-PG | 15 | 7.5 | 3.75 | 1.88 | 0.94 | 0 |
| **9** | 16:0-18:1 D5 PG | 15 | 7.5 | 3.75 | 1.88 | 0.94 | 0 |
| **10** | 15:0-18:1-d7-PS | 20 | 10 | 5 | 2.5 | 1.25 | 0 |
| **11** | 14:0 Cardiolipin | 10 | 5 | 2.5 | 1.25 | 0.63 | 0 |
| **12** | 18:1-d9 SM | 2.5 | 1.25 | 0.63 | 0.31 | 0.16 | 0 |
| **13** | C15 Ceramide-d7 (d18:1-d7/15:0) | 5 | 2.5 | 1.25 | 0.63 | 0.31 | 0 |
| **14** | 18:1-d7 MG | 25 | 12.5 | 6.25 | 3.13 | 1.56 | 0 |
| **15** | 15:0-18:1-d7 DG | 3 | 1.5 | 0.75 | 0.38 | 0.19 | 0 |
| **16** | 15:0-18:1-d7-15:0 TG | 1.5 | 0.75 | 0.38 | 0.19 | 0.09 | 0 |
| **17 (22)** | 17:0-17:1-17:0 D5 TG | 1.5 | 0.75 | 0.38 | 0.19 | 0.09 | 0 |
| **18 (23)** | 1,3-17:0 D5 DG | 2.5 | 1.25 | 0.63 | 0.31 | 0.16 | 0 |
| **19** | cholesterol-d7 | 22.5 | 11.25 | 5.63 | 2.81 | 1.41 | 0 |
| **20** | 15:0 cholesteryl-d7 ester | 15 | 7.5 | 3.75 | 1.88 | 0.94 | 0 |
| **21** | Cardiolipin Mix I | 15 | 7.5 | 3.75 | 1.88 | 0.94 | 0 |
| **22** | d5-TG ISTD Mix I | 2.5 | 1.25 | 0.63 | 0.31 | 0.16 | 0 |
| **23** | d5-DG ISTD Mix I | 7.5 | 3.75 | 1.88 | 0.94 | 0.47 | 0 |
| **24** | Cer/Sph Mixture I | 7.5 | 3.75 | 1.88 | 0.94 | 0.47 | 0 |

**Table A.2.** Optimal Peak detection parameter setting in mzMine and XCMS

| Module | mzMine | | XCMS | |
|---|---|---|---|---|
| Mass detection | Noise level | 1.0·$10^5$ | Noise | 1.0·$10^5$ |
| EIC building | Min group size in # of scans | 4 | firstBaselineCheck | TRUE |
| | Group intensity threshold | 1.5·$10^5$ | | |
| | Min highest intensity | 2.0·$10^5$ | prefilter | c(1, 2.0·$10^5$) |
| | m/z tolerance | 7 ppm | mzTolerance | 7 ppm |
| Chromatogram peak picking | Algorithm | Wavelets(ADAP) | method | centWave |
| | m/z center calculation | Auto | mzCenterFun | wMean |
| | S/N threshold | 10 | snthresh | 10 |
| | S/N estimator | Intensity window SN | integrate | 2 |
| | Peak duration range | 0.04-0.5 minutes | peakwidth | 2.4-30 seconds |
| | RT wavelet range | 0.004-0.05minutes | | |
| | min feature height | 2.0·$10^5$ | | |
| | coefficient/area threshold | 100 | | |

In the mass detection module, the *noise level* for mzMine and XCMS are both set to 1.0·$10^5$. In the EIC building module, the *firstBaselineCheck* in XCMS is set as True. In this algorithm, the *sequential points above baseline* parameter is similar to mzMine's *Min group size in # of scans*, while the *Baseline estimation* is similar to mzMine's *Group intensity threshold*. The *prefilter* and *mzTolerance* parameters in XCMS are similar to mzMine's *Min highest intensity* and *mz tolerance*, which are set as 2.0·$10^5$ and 7 ppm separately. The above parameters are set according to the visualization of the peaks.

In the Chromatogram peak picking module, *Wavelets (ADAP)* and *centWave* algorithms are chosen for mzMine and XCMS respectively. Both of them are based on the wavelet chromatographic peak detection algorithm. The *calculation of mz center* is selected as *Auto* (automatic $\log_{10}$ weighted approach) for mzMine and *wMean* (intensity weighted mean of the peak's *mz* values) for XCMS. The *S/N estimator* is selected as Intensity window SN in mzMine, which is similar to XCMS's integrate 2 parameter, this parameter means peak limits are found through descent on the real data. mzMine's *Peak duration range* is set as 0.04-0.5 minutes. Correspondingly, the XCMS's *peakwidth* is set as 2.4-30 seconds. mzMine contains other 3 unique parameters: *RT wavelet range*, *min* *feature height* and *coefficient/area threshold*, which are set as 0.004-0.05 minutes, $2.0 \cdot 10^5$ and 100 respectively. Among them, *RT wavelet range* indicates the range of wavelet scales used to build a matrix of coefficients. Scales are expressed as RT values (minutes) and correspond to the range of wavelet scales that will be applied to the chromatogram; while *min feature height* indicates the minimum height of the feature considered as detected; *coefficient/area threshold* is the best coefficient found by taking the inner product of the wavelet at the best scale and the peak, and then dividing by the area under the peak.

Progenesis is commercial software, of which the algorithm of the peak detection is not available. For this reason the parameters of Progenesis are not shown here.

**Table A.3.** Grouping parameters

| **mzMine** | | **XCMS** | |
|---|---|---|---|
| **Algorithm** | Join aligner | Algorithm | PeakDensityParam |
| **m/z tolerance** | 0.02 Da | binSize | 0.02 Da |
| **RT tolerance** | 0.1 abs min | Bw | 6s |
| **Weight for RT** | 0.5 | minFraction | 0.5 |
| **Weight for m/z** | 0.5 | minSamples | 1 |

The grouping parameters for mzMine and XCMS can be accessed via Table S5. mzMine's *mz tolerance* (0.02 Da) and *RT tolerance* (0.1 absolute minutes) are comparable to XCMS's *binSize* (0.02 Da) and *Bw* (6 Second). mzMine's *Weight for RT, Weight for mz* are unique parameters. In contrast to XCMS's unique parameters: *minFraction* (minimum fraction of samples in at least one sample group in which the peaks have to be present to be considered as a peak group), *minSamples* (minimum number of samples in at least one sample group in which the peaks have to be detected to be considered a peak group).

**Table A.4.** Isotope filtration parameters

| mzMine | | XCMS | |
|---|---|---|---|
| **Ionization Polarity** | positive | polarity | positive |
| **FWHM sigma** | 6 | sigma | 6 |
| **FWHM percentage** | 60% | perfwhm | 0.6 |
| **Isotopes max. charge** | 1 | maxcharge | 1 |
| **Isotopes max. per cluster** | 4 | maxiso | 4 |
| **Isotopes mass tolerance** | 5ppm or 0.01 mz | ppm | 5 |
| | | mzabs | 0.01 |
| **Correlation threshold** | 0.75 | cor_eic_th | 0.75 |
| **Correlation p-value** | 0.05 | pval | 0.05 |

The table above shows the isotope filtration parameters for mzMine and XCMS. Both of them use the CAMERA's algorithm[4] for isotope filtration.

**Table A.5.** Internal standards quantification

| Label | Compound | Adduct | mz | XCMS ·$10^7$ | mzMine·$10^7$ | Progenesis·$10^7$ |
|---|---|---|---|---|---|---|
| A | 20:0-20:1-20:0 D5 TG | M+Na | 1000.929 | 26.206 | 26.095 | 12.425 |
| B | 18:1-d7 MG | M+H | 364.344 | 3.214 | 3.237 | 0.089 |
| C | 17:0-17:1-17:0 D5 TG | 2M+Na | 1726.587 | 1.980 | 2.934 | 23.802 |

In the table, the abundances calculated by "area" from XCMS and mzMine are used, while the abundance called "raw abundance" from Progenesis is used. These three Compounds share the same abundance trend in XCMS and mzMine (i.e. 26.206, 3.214 and 1.980 ·107 for compound A, B and C respectively in XCMS), while quite different in Progenesis (i.e. 12.425, 0.089 and 23.802 ·107 for compound A, B and C respectively in Progenesis).
